# DynamicDemiLog: A Single Sketch for Ultrafast Similarity, Frequency, and Cardinality Estimation

**DOI:** 10.64898/2026.06.12.731986

**Authors:** Brian Bushnell

## Abstract

Probabilistic cardinality estimators (HyperLogLog), similarity sketches (MinHash), and frequency estimators (Count-Min Sketch) are fundamental approximate data structures that each target one primary problem. We present DynamicDemiLog (DDL), a sketch that unifies cardinality estimation, set similarity, containment, element frequency and composition in one tiny data structure built from a single pass over the input stream. Using an inverted index over 200,687 RefSeq sketches (159,567 organisms), DDL performs all-to-all sketch similarity comparison of the full database in 30 seconds (128 threads, indexed) — over 375× faster per query than Mash’s brute-force all-to-all comparison of 91,282 sketches, or 31× faster without the index, at double the sketch resolution. DDL extends the LogLog register with a *mantissa*: each register stores a floating-point-encoded hash value consisting of an integer exponent (the leading-zero count) and a fractional mantissa (the sub-leading-zero bits), rather than the integer leading-zero count alone. This preserves enough hash information for meaningful register-by-register comparison — a property that standard 6-bit registers lack — while improving on LogLog’s cardinality estimation machinery, including DynamicLogLog’s early exit mask for high-throughput streaming.

With a default 10 mantissa bits (16-bit registers, 2,048 buckets, 4 KB), DDL achieves a per-register false-match rate of 0.018% on unrelated random same-size sets (compared to 17.0% for LL6, a basic HyperLogLog implementation), enabling Weighted Kmer Identity (WKID), Average Nucleotide Identity (ANI), containment, and completeness estimation from register comparison alone. A 16-bit per-register observation counter provides element frequency information at trivial additional computation cost, and an additional byte tracks element composition (GC content, for biological data). Furthermore, DDL’s high-specificity registers enable an inverted index structure (DDLIndex) that answers similarity queries against a database of *N* sketches in O(*B* + *M*) time, where *M* is the number of matching index entries, compared to O(*N* ×*B*) for pairwise comparison.

DDL achieves a 930× reduction in false register matches compared to LL6 (Section 11.1), accurately estimates ANI between full and partial genomes down to approximately 79% identity (at k=25, B=2,048), and maintains near-zero spurious similarity on unrelated inputs — all at similar construction speed to LL6, and 3.5× faster than SetSketch.

## 1. Introduction

Streaming algorithms that summarize large datasets in bounded space are foundational tools across computer science. Four problems recur across nearly every domain that processes high-volume data streams:

1. **Cardinality estimation** — how many distinct elements are in the stream? Used in network monitoring (distinct IP flows), database optimization (query planning), and memory requirement estimation.
2. **Set similarity** — how similar are two streams? Used in document deduplication, malware clustering, plagiarism detection, near-duplicate and containment detection.
3. **Frequency estimation** — how often does each element appear? Used in anomaly detection, heavy-hitter identification, and workload characterization.
4. **Composition analysis** — what type of elements occur at which frequency? Used in contamination detection, efficiency profiling, and proposed for data-agnostic set signatures. No well-known bounded-space algorithm addresses this problem; existing tools such as Jellyfish (Marçais and Kingsford 2011), KMC (Kokot, Długosz, and Deorowicz 2017), and KAT (Mapleson et al. 2017) require memory proportional to the number of distinct elements.

The first three problems have well-established solutions: HyperLogLog (Flajolet et al. 2007) for cardinality, MinHash (Broder 1997) for Jaccard similarity, and Count-Min Sketch (Cormode and Muthukrishnan 2005) for frequency estimation. Each is elegant, efficient, and widely deployed. But each is also *isolated*: a system that needs all three must maintain multiple separate data structures, process each element multiple times, and use additional memory. Furthermore, Count-Min Sketches decline in accuracy with increased cardinality, and thus require memory proportional to cardinality for high accuracy — like exact counting, but with a lower constant.

This fragmentation is not merely an engineering inconvenience — it reflects a deeper limitation in how these algorithms use hash information. HyperLogLog stores only the Number of Leading Zeros (NLZ) in each register, discarding the sub-NLZ bits entirely. This is sufficient for cardinality (the NLZ encodes the approximate magnitude of the maximum hash per bucket) but too coarse for comparison: two registers with the same integer NLZ are likely to contain entirely different elements, making register-level comparison meaningless. MinHash stores the full minimum hash value per bucket, preserving comparison ability and cardinality estimation, but at the cost of substantially more memory and the computational complexity of maintaining a sorted set during construction. Extending this data structure to support composition and frequency further reduces speed and increases complexity (though BBSketch (Bushnell 2016) does maintain counts for MinHash). Count-Min Sketch achieves collision resilience through multi-hashing while discarding all information about individual elements, thus allowing cardinality estimation, frequency analysis, and occurrence counts of any specific element, but composition tracking becomes impossible.

DynamicDemiLog (DDL) — “demi” because its register width is half fixed (the mantissa) and half logarithmic (the NLZ exponent), no longer a pure LogLog — bridges this gap with a single architectural change: it stores each register as a *floating-point-encoded hash value*, consisting of an integer exponent (the NLZ) and a fractional mantissa (the bits immediately following the leading one). This is analogous to IEEE 754 floating-point representation, where the exponent captures order of magnitude and the mantissa captures precision within that order.

This change has three consequences:

- **Cardinality estimation is improved.** The exponent (NLZ) feeds the same estimation pipeline as HyperLogLog and DynamicLogLog — DLC, Hybrid, and correction-factor-corrected (CF-corrected) Mean all work unchanged. The mantissa additionally enables MeanM, a novel estimation method that reconstructs approximate hash magnitudes from the stored (exponent, mantissa) encoding and estimates cardinality from their arithmetic mean — enabling increased accuracy per bucket, without requiring the precomputed terminal correction factors that integer-NLZ estimators need (Section 4.1).
- **Register comparison becomes meaningful.** With *m* mantissa bits, the probability of two unrelated elements producing the same stored value (given the same NLZ tier) drops from 1 (at *m* = 0, i.e., standard HLL) to approximately 1/2^*m*. At *m* = 10, this is ∼1/1,024 — specific enough that matching registers strongly imply shared elements. This enables WKID, ANI, containment, and completeness estimation from register comparison alone.
- **Frequency tracking is free.** Leveraging DynamicLogLog’s *dynamic early exit* mask (eeMask, Section 3.2), DDL rejects the vast majority of elements before any memory access. Empirically (measured by counting taken/not-taken branches), the rejection rate exceeds 99.9% for datasets with high cardinality-to-bucket ratios (e.g., eukaryotic genomes or raw read sets at *B* = 2,048) and exceeds 97% even for small genomes such as *E. coli* (∼4.6M unique k-mers at *B* = 2,048). The rejection rate depends on both *N* and *B* independently, not simply *N* /*B*, because all *B* buckets must advance past the minimum tier — a coupon-collector effect that requires superlinearly more elements per bucket as *B* grows. The fewer than 0.1–3% of elements that survive this filter and actually update a register can concurrently update per-register frequency counters and GC-content trackers at negligible cost. This is why “Dynamic” appears in the name: the dynamic early exit makes auxiliary per-register tracking essentially free.

Beyond pairwise comparison, DDL’s high-specificity registers enable an *inverted index* (DDLIndex, Section 6) that maps each (bucket, value) pair to a list of sketch IDs. With 16-bit registers (65,536 possible values per bucket), the index is both memory-efficient and specific enough for low false-positive rates, enabling ∼40× speedup over brute-force pairwise comparison for database search (96.3% index efficiency; Section 11.7).

### Related work

Several recent works have extended LogLog-type sketches beyond cardinality estimation. HyperMinHash (Yu and Weber 2022) independently introduced the same floating-point register encoding. HyperMinHash typically uses 10-bit registers (6-bit exponent + 4-bit mantissa) and provides cardinality, Jaccard, and intersection estimation. DDL uses wider 16-bit registers, which provides three practical advantages: (1) the default 10-bit mantissa gives a theoretical per-register false match rate ∼64× lower than HyperMinHash’s proposed 4-bit mantissa (1,024 vs. 16 intra-tier values; no empirical comparison is possible due to the lack of a public HyperMinHash implementation), enabling reliable pairwise comparison; (2) the 65,536-value register space enables an efficient inverted index (Section 6); and (3) 16-bit registers align to hardware word boundaries and enable 16-way SIMD comparison via AVX-256 or NEON. DDL additionally provides capabilities HyperMinHash lacks: directional comparison (lower/equal/higher per bucket, enabling containment and completeness), per-register frequency counting and composition tracking, and a dynamic early exit mask inherited from DynamicLogLog.

SetSketch (Ertl 2021) replaces the leading-zero count with a truncated logarithm of configurable base, providing cardinality and Jaccard estimation from a single sketch with maximum-likelihood estimators. With base *b* = 1.001 and 2-byte registers (*q* = 65,534), SetSketch uses the same memory as DDL but encodes register values as a 16-bit fine-grained exponent with no separate mantissa. Empirically, this encoding produces 40% more false register matches than DDL on unrelated same-size random input (0.52 vs. 0.37 per pair; Section 11.1), because the logarithmic encoding clusters values where the hash magnitude is dense, whereas DDL’s 10+-bit mantissa provides effectively random discrimination within each exponent tier. SetSketch’s per-element logarithm computation also makes it 3.5× slower to construct than DDL (Section 11.4). SetSketch does not provide frequency tracking, composition tracking, directional comparison, or an inverted index; its collision rate analysis focused on Jaccard RMSE rather than per-register false match rates.

Dashing (Baker and Langmead 2019) and Dashing2 (Baker and Langmead 2023) apply HLL to genomic ANI estimation via joint maximum likelihood estimation (JMLE). Dashing2 switched from HLL to SetSketch internally. Neither provides per-register directional comparison, frequency counters, or inverted index support.

DDL’s novelty lies not in the floating-point register encoding itself — which HyperMinHash introduced in 2017 (arXiv preprint; published 2022 (Yu and Weber 2022)), with BBTools’ LogLog16 independently implementing a similar encoding in 2020 — but in the *combination* of capabilities built on that encoding: directional register comparison, frequency counting, composition tracking, inverted index, SIMD-friendly 16-bit registers, and dynamic early exit — unified in a single data structure, along with integrated support for genomic sequence data and a prebuilt index of all 159,567 organisms in RefSeq (200,687 sketches, with separate sketches for organelles).

DDL is implemented in Java as part of the BBTools suite (Bushnell 2014) and is available as `DynamicDemiLog.java` in the `cardinality` package, usable via `loglog.sh` for k-mer frequency and composition histograms. The comparison tool is `DDLCompare.java` in the `ddl` package, usable via `ddlcompare.sh` or `quickclade.sh`. The RefSeq index is available at https://github.com/bbushnell/BBTools/releases/ and is also accessible without download via remote sketch queries using QuickClade (Bushnell, Schulz, and Villada 2026).

## 2. Background

### 2.1 LogLog-Family Cardinality Estimation

All LogLog-family estimators share a common structure. Given a hash function mapping elements to uniformly distributed integers in [0, 2^*L*), the hash is split into a *bucket selector* (the lowest *k* bits, selecting one of *B* = 2^*k* buckets) and a *rank* (the Number of Leading Zeros in the remaining bits). Each bucket stores the maximum NLZ observed. The collection of per-bucket maximum NLZ values encodes the approximate cardinality of the stream.

HyperLogLog (Flajolet et al. 2007) estimates cardinality as:

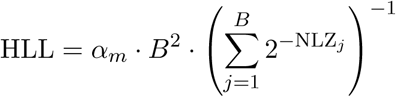

where *α_m_* ≈ 0.7213*/*(1 + 1.079*/B*) is a bias-correction constant. At low cardinality, Linear Counting (LC = *B ·* ln(*B/V*), where *V* is the number of empty buckets) is used instead. The transition between LC and HLL produces a characteristic error spike — see (Bushnell 2026b) for detailed analysis.

DynamicLogLog (DLL) (Bushnell 2026b) describes a shared exponent (`minZeros`) across all buckets, storing only the relative NLZ per bucket. This reduces register width from 6 bits to 4, enables an early exit mask that rejects 97–99.9%+ of elements depending on cardinality (Section 3.2), and introduces Dynamic Linear Counting (DLC) — a tier-aware extension of LC that provides accurate estimates across the full cardinality range without a transition spike.

### 2.2 MinHash Similarity Estimation

MinHash (Broder 1997) estimates the Jaccard similarity of two sets by hashing all elements and retaining the *k* minimum hash values per set. The fraction of matching minimums estimates the Jaccard index:

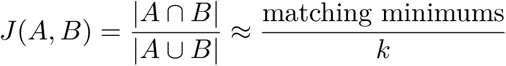

Mash (Ondov et al. 2016) applies MinHash to genomic k-mers, estimating Average Nucleotide Identity (ANI) from the Jaccard index via a mutation model.

MinHash stores full hash values (typically 64 bits each), preserving the information needed for comparison and cardinality estimation but requiring a sorted set of large numbers, which is inefficient in both time and space.

BBSketch (Bushnell 2016), introduced shortly after Mash in 2016, extends MinHash with dual k-mer lengths (k=32 and k=24 simultaneously), per-hash occurrence counting, estimation of genomic completeness and contamination, real-time gene-calling and protein-space sketching, key indexing, variable-length sketch comparison, SSU embedding for alignment ANI, entropy filtering, and a blacklist for taxonomically uninformative k-mers. BBSketch’s extended feature set comes at the cost of higher computational and memory overhead relative to basic MinHash.

For exact frequency analysis, KmerCountExact and BBNorm (Bushnell 2014) provide exact and Count-Min Sketch representations of k-mer sets, respectively. Both support k-mer frequency histograms, cardinality counting, and genome size estimation from histogram peaks. KmerCountExact additionally supports compositional (GC) reporting, but at memory cost proportional to the number of distinct k-mers — impractical for complex metagenomic libraries.

### 2.3 Notation

**Table 1:**
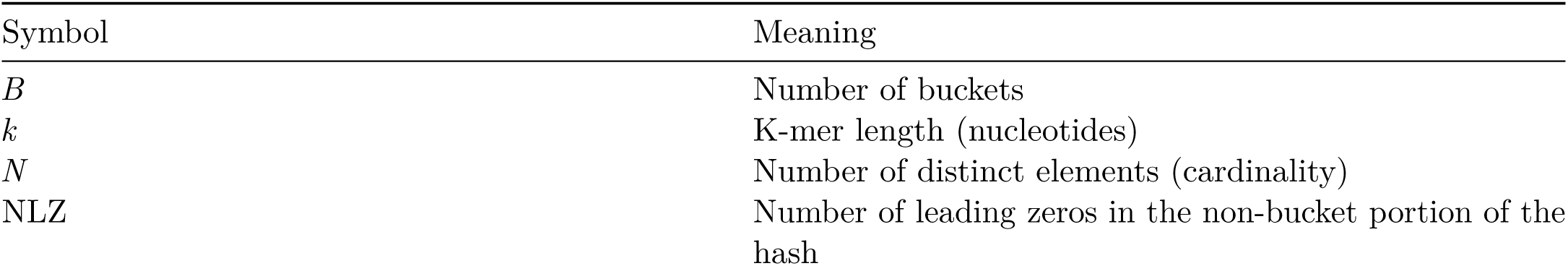
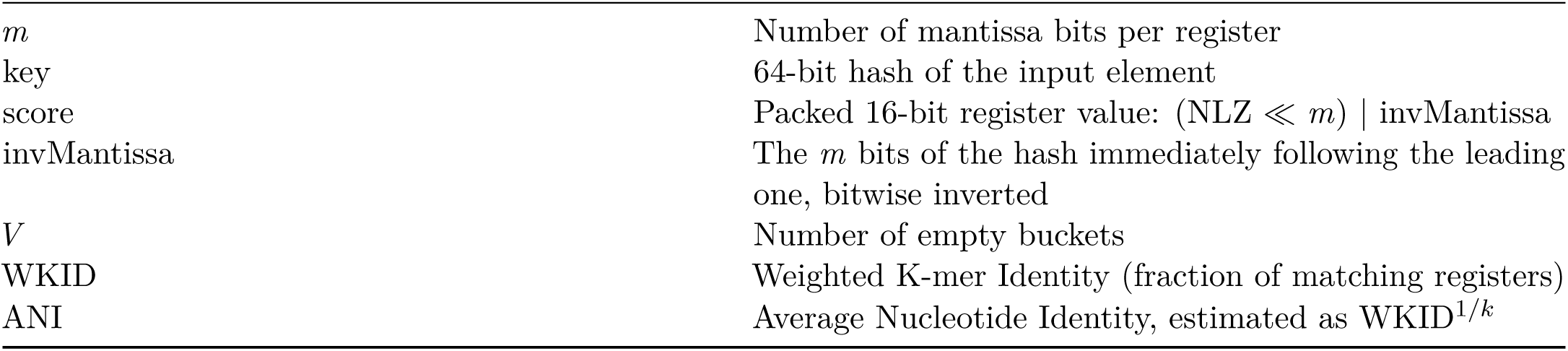
Notation used throughout this paper.

## 3. DDL Architecture

### 3.1 Floating-Point Register Encoding

DDL stores each register as a 16-bit unsigned integer encoding a floating-point hash value:

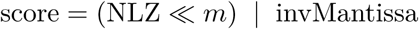

where NLZ is the Number of Leading Zeros (the exponent) and invMantissa is the inverted mantissa — the *m* bits immediately following the leading one in the hash value, bitwise inverted.

In the default configuration, the exponent occupies the upper 6 bits (supporting NLZ values 0–62, generally sufficient for 64-bit hashes), and the mantissa occupies the lower *m* = 10 bits (Figure 1). The exponent/mantissa split is configurable at runtime (Section 8.2). A stored value of 0 indicates an empty register; the minimum non-empty stored value is exponent 1 (indicating NLZ=0), mantissa 0.

**Figure 1:**
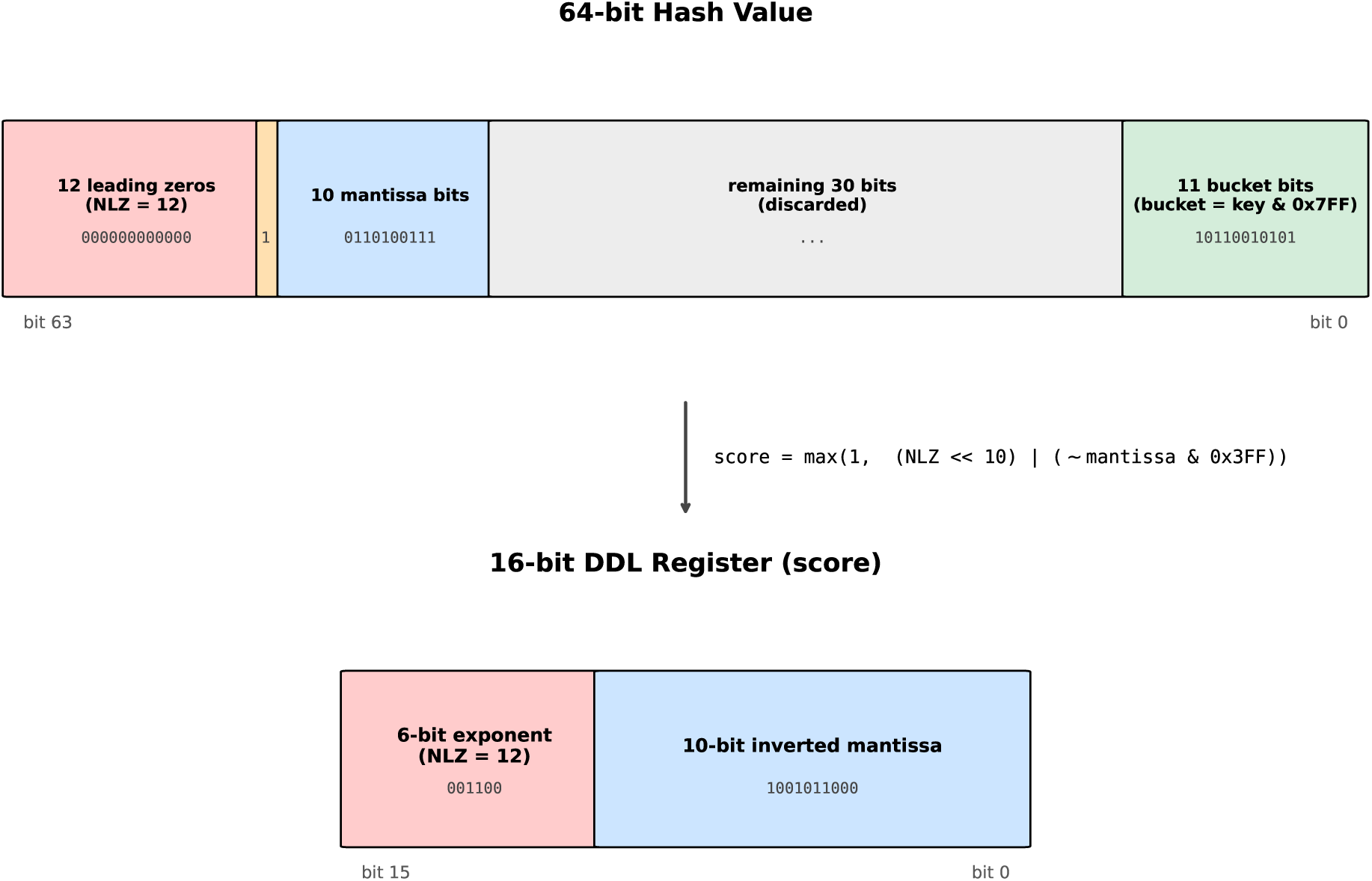
DDL register encoding (default 6e+10m configuration). A 64-bit hash is decomposed into bucket-selector bits (lower 11), leading zeros (NLZ, the exponent), a leading one (discarded), and mantissa bits (the *m* bits after the leading one). The 16-bit register stores the NLZ in the upper exponent bits and the inverted mantissa in the lower *m* bits, forming a single unsigned score that increases monotonically with hash quality.

### Inverted mantissa convention

The mantissa bits are *inverted* (bitwise NOT) before storage. This encoding has a critical property: for two elements hashing to the same NLZ tier, the one with a *smaller* raw hash value (a rarer event) produces a *higher* inverted mantissa. Combined with the NLZ exponent in the upper bits, this means the stored score increases monotonically with hash “quality” — higher NLZ is always better, and within the same NLZ, higher inverted mantissa indicates a rarer hash.

This makes the register update condition a single unsigned comparison:

`if (newScore > oldScore) { register = newScore; }`

The same comparison that updates cardinality information also preserves the most informative hash for similarity comparison. No separate logic is needed to maintain the mantissa.

### 3.2 Hash Processing

The complete hash-and-store operation for DDL:

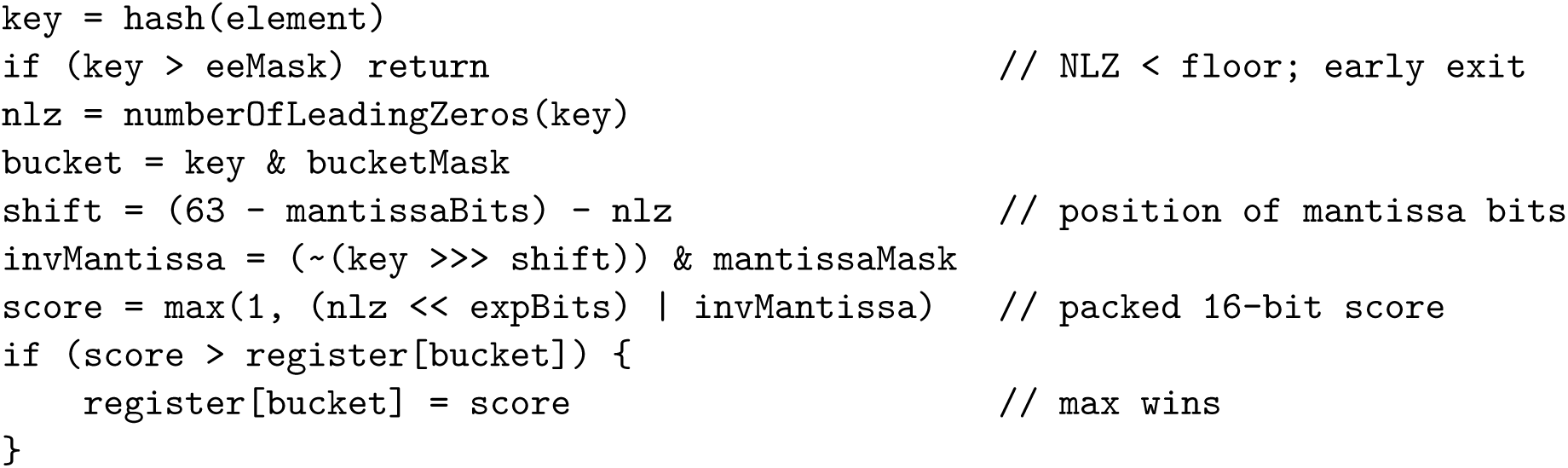

The early exit mask (eeMask) is inherited directly from DynamicLogLog. It tracks the global NLZ floor — the minimum NLZ across all registers — and rejects any hash whose NLZ falls below this floor, since it cannot possibly update any register. At high cardinality, the vast majority of elements are rejected before any register access.

Unlike DynamicLogLog’s DLL4 variant, DDL stores *absolute* NLZ values rather than relative ones. This retains full idempotency — important for sketch comparison, where a sketch’s value must not depend on insertion order, unlike DLL4’s relative encoding where values shift as the global floor rises. This simplifies merge operations — two DDL sketches can be merged by taking the per-register maximum directly, retaining the composition and count values of the winner, or summing the counts in a tie.

### 3.3 Memory Layout

With 16-bit registers, DDL stores 2,048 buckets in 4,096 bytes (4 KB). Compare:

**Table 2:** Memory per sketch at 2,048 buckets.

| Sketch | Bits/register | 2,048 registers | Information content |
| --- | --- | --- | --- |
| HLL (6-bit) | 6 | 1,536 B | NLZ only |
| DLL4 (4-bit) | 4 | 1,024 B | Relative NLZ only |
| DDL (16-bit) | 16 | 4,096 B | NLZ + mantissa |
| MinHash (64-bit) | 64 | 16,384 B | Full hash value |

DDL uses 2.67× the memory of packed 6-bit HLL and 4× the memory of DLL4, but provides similarity and frequency estimation that neither offers. In practice, HLL implementations often store 6-bit values in 8-bit registers (2 KB at 2,048 buckets), making DDL’s practical overhead 2× rather than 2.7×. Compared to MinHash, DDL uses 1/4 the memory (or 5/8 including the count and composition) while additionally allowing storage in a simple array rather than the set and/or heap needed for efficient MinHash construction — the intermediate and final forms of DDLs are identical. This contrasts with the intermediate LongHeapSet used by BBSketch, roughly 3x the size of the final sketch.

## 4. Cardinality Estimation

DDL supports all of DynamicLogLog’s cardinality estimators. The exponent portion of each register is functionally identical to DLL’s stored NLZ, so DLC (Dynamic Linear Counting), the Logarithmic Hybrid Blend, and correction-factor-corrected (CF-corrected) Mean all work unchanged — the mantissa is simply ignored for these estimation methods.

### 4.1 MeanM: Correction-Factor-Free Estimation

The mantissa enables a fundamentally different approach to cardinality estimation. Standard LogLog estimators work only with integer NLZ values, which coarsely quantize the hash magnitude. This quantization introduces systematic bias that varies non-trivially with cardinality and must be compensated by empirically-derived correction factor tables that asymptotically approach a constant — a practical nuisance and a source of residual error. Due to tracking a continuous sub-NLZ fraction, DDL’s asymptotic constant is within 1/(2^mantissa bits) of 1.0; thus it does not need a correction factor above very low cardinality where some buckets are empty.

DDL’s floating-point registers store enough hash information to reconstruct an *approximate hash magnitude* from each register. The `restore` operation inverts the floating-point encoding:

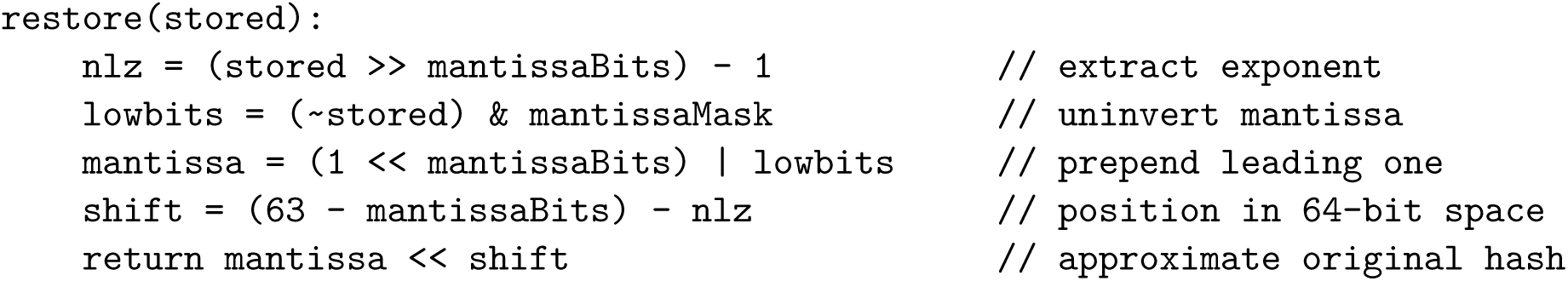

This yields a value with *m*+1 bits of precision (1 implicit + *m* mantissa) positioned by the NLZ exponent — exactly analogous to converting a floating-point number back to its integer representation. With integer NLZ alone, each tier produces a single magnitude (2^(63-NLZ)); with the mantissa, each tier is subdivided into 2^*m* distinct magnitudes.

The arithmetic mean of restored values directly estimates cardinality without correction factors:

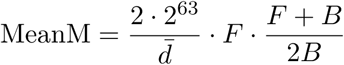

where 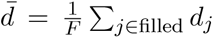 is the arithmetic mean of restored hash values across the *F* filled buckets, and the (*F* + *B*)*/*(2*B*) term is a simple linear correction for empty buckets. The core estimate 2 · 2^63^*/d̅* follows from the observation that if the minimum hash per bucket has average magnitude *d̅*, then approximately 2^64^/*d̅* elements were seen — the same logic as HyperLogLog, but with more precise per-register values.

### MeanM approximates the maximum likelihood estimator

Each restored value *d_j_* approximates the minimum hash in bucket *j*. Define 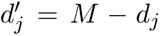, where *M* = 2^64^; since the minimum of *n* uniform random variables on [0*, M*) and *M* minus their maximum have the same distribution, 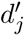 follows the CDF of the maximum of *n* = *N/B* independent Uniform(0, M) values:

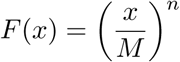

The log-likelihood of observing *B* independent maxima 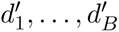 from this distribution is 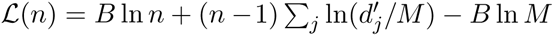. Setting *∂*L*/∂n* = 0 yields the maximum likelihood estimator:

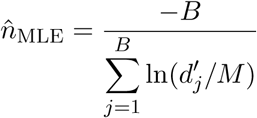

For *d_j_*≪ *M* (high cardinality, where most hash magnitudes are small relative to the full range): 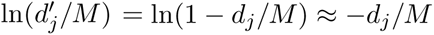, so:

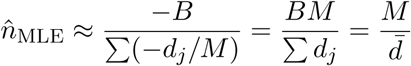

and *N̂* = *n̂* · *B* = *BM/d̅* — which is exactly MeanM (up to the empty-bucket correction). MeanM is therefore a first-order approximation to the MLE that becomes exact as cardinality grows and *d_j_/M* → 0.

The mantissa’s contribution to estimation accuracy is modest but measurable. Inte<u>g</u>er-NLZ estimators are limited by quantization of the hash magnitude to an asymptotic relative error of 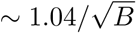 (the HyperLogLog const<u>an</u>t), while MeanM’s first-order equivalence to the MLE brings DDL within 1*/*2*^m^* of the Cramér–Rao bound of 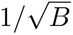 — at *B* = 2,048, 2.21% versus 2.30%, a 3.8% reduction in relative error. Since no unbiased estimator can improve on 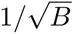, MeanM is within a quantization term of optimal: accuracy gains beyond this point require more buckets, not better estimation. The mantissa’s larger benefit is structural — correction-factor-free estimation across the full cardinality range, and the lower collision rate.

### 4.2 Production Estimator

DDL’s production estimator (hybridDDL) uses a three-zone blend:

1. Below 0.2*B*: pure Linear Counting (LCmin)
2. 0.2*B* to 7.5*B*: log-interpolated blend from LCmin to CF-corrected MeanM
3. Above 7.5*B*: pure CF-corrected MeanM

At low cardinality, Linear Counting is most accurate (it uses empty-bucket counts, which are maximally informative when many buckets are empty). At high cardinality, the mantissa-based MeanM estimator dominates, providing a CF-free arithmetic mean that eliminates the quantization bias inherent in integer-NLZ estimation. Since the true cardinality is not known, for the purpose of blending, it is estimated from the mantissa-free, CF-free DLC formula (Bushnell 2026b).

Residual correction factors from DynamicLogLog’s CF infrastructure can optionally be applied but contribute only at low cardinality — the mantissa has already eliminated the dominant quantization bias.

## 5. Set Comparison

### 5.1 Register-Level Comparison

Given two DDL sketches A and B with the same hash seed and bucket count, every bucket can be classified by comparing stored scores:

- **Lower:** A’s register < B’s register (B has a rarer hash in this bucket)
- **Equal:** A’s register = B’s register (likely the same element won both)
- **Higher:** A’s register > B’s register (A has a rarer hash in this bucket)
- **Both empty:** both registers are 0

The comparison is branchless (shown here for the three-way case; the implementation additionally tracks both-empty pairs via a separate pre-check):

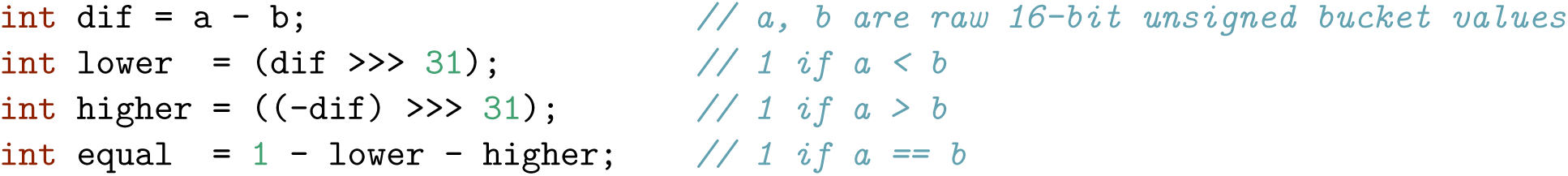

### 5.2 Why the Mantissa Makes Comparison Meaningful

With integer NLZ alone (standard HLL), two registers from unrelated streams that happen to land in the same NLZ tier are guaranteed to “match” — every NLZ=5 register looks the same. Empirically, 17% of LL6 register pairs produce spurious matches on same-sized unrelated random sets (Section 11.1) — too high for meaningful comparison. This makes HLL register comparison insufficient for similarity estimation.

With *m* mantissa bits, a tier is subdivided into 2^*m* distinct stored values. Two unrelated elements in the same tier produce a matching stored value with probability approximately 1/2^*m*. At *m* = 10, this is approximately ∼1/1,024 — specific enough that a matching register strongly implies a shared element.

### Expected false matches

Empirically, DDL (6e+10m) averages 0.37 false register matches per unrelated pair — a 930× reduction from LL6’s 347.5, explained by the default mantissa’s ∼1/1,024 intra-tier discrimination (Section 11.1).

### 5.3 Similarity Metrics

#### WKID (Weighted K-mer IDentity)

The primary similarity metric, denoted “Weighted” to differentiate it from raw K-mer IDentity (KID), the simple fraction of shared/total values. WKID weights by genome size to project “What would the KID be if the genomes were the same size”, such that comparing half of a set with the full set will yield KID of 0.5, but WKID of 1.0.

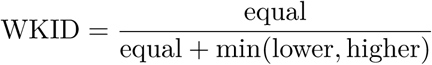

The min(lower, higher) in the denominator selects the mismatch count from the *smaller* set’s perspective. When two sets of unequal size are compared, the larger set has more unique elements and therefore more buckets where it “wins” — using min prevents this asymmetry from inflating the denominator. This is analogous to BBSketch’s minDivisor approach.

When the divisor is small (below 6), it is padded to prevent inflated scores from random bucket collisions between pairs with highly dissimilar cardinality.

#### ANI (Average Nucleotide Identity)

For k-mer-based sketches, WKID measures the fraction of registers sharing the same winning k-mer — a quantity that tracks the k-mer survival probability under mutation. At per-nucleotide identity ANI, a k-mer of length *k* survives intact with probability approximately ANI^*k*, so WKID ≈ ANI^*k*. ANI is therefore estimated as:

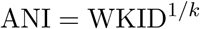

This uses the same uniform random SNP-based mutation model used by BBSketch, Mash, and other k-mer-based tools. However, unlike Mash — which uses raw Jaccard (equivalent to KID) as its base — DDL and BBSketch use WKID, enabling correct ANI estimation between genome fragments and full genomes: comparing reads from half of *E. coli* against the full genome yields DDL ANI = 100% and completeness = 50%, whereas Mash reports ANI ≈ 98.7% on identical sequence because its KID-based formula interprets partial coverage as mutation. The estimate is accurate down to approximately 79% ANI at *B* = 2,048 and *k* = 25 (5e+11m), based on the reliability floor defined in Section 8.3; below this, too few k-mers survive mutation for reliable estimation.

### Containment

The fraction of A’s content shared with B, analogous to Broder’s containment metric (Broder 1997):

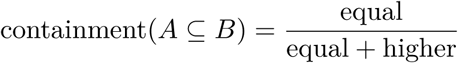

From A’s perspective, *higher* counts the buckets where A has a rarer hash than B — these represent elements unique to A. If A is a subset of B, A should have few unique elements and containment approaches 1.

### Completeness

The relative size of A compared to B:

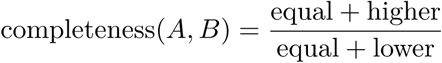

This estimates the fraction of B’s content present in A, clamped to [0, 1]. If A is a pure subset of B (containing half of B’s k-mers with no unique content), then Higher ≈ 0 and Equal ≈ Lower, yielding completeness ≈ 0.5 — the correct fraction. Crucially, this differs from a simple cardinality ratio |A|/|B|: a set A with high cardinality but little overlap with B would have many Higher buckets (unique to A) but few Equal buckets (shared), yielding low completeness despite high cardinality. Thus completeness measures *shared content fraction*, not relative size.

Empirical validation: synthetic 10× reads from a 50% fragment of *E. coli* K-12 (generated by `mutate.sh fraction=0.5` and `randomreadsmg.sh`) compared against the full genome yield DDL completeness of 0.5005, CompareSketch completeness of 49.57%, and DDL WKID of 1.0000 (ANI = 100%) — correctly identifying the sequence as identical but partial.

## 6. Inverted Index

### 6.1 DDLIndex Structure

DDL’s 16-bit registers create a structured value space of 65,536 possible stored values per bucket. This enables an *inverted index* — a data structure that maps each (bucket, value) pair to a list of sketch IDs:

`index[bucket][value] → list of sketch IDs with that value in that bucket`

The index is a 2D array of *B* × 65,536 cells. In practice, register values cluster in the NLZ tiers corresponding to genomic cardinalities in the database, leaving approximately 73% of cells null — the value space is sparse despite containing hundreds of thousands of sketches.

### 6.2 Query Processing

To find all sketches similar to a query, DDLIndex scans the query’s *B* registers and looks up each (bucket, value) pair in the index and increments counters of reference sketches in a sparse set:

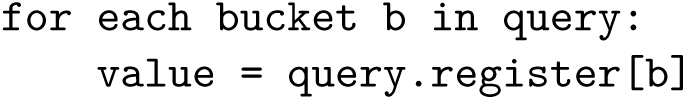
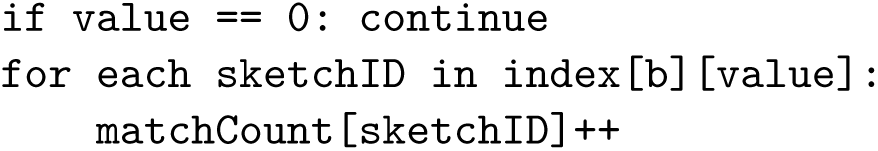

A sketch with many matching registers has high WKID with the query. The query time is O(*B* + *M*), where *M* is the total number of matching index entries — independent of the number of sketches in the database when the index is sparse. This is in contrast to exhaustive pairwise comparison, which requires O(*N* × *B*) time for *N* database entries. Subsequently, only references exceeding a minimum match threshold are directly compared using the full pairwise comparison (Section 5.1).

### ANI from index matches

The index directly provides a match count per sketch. ANI can be computed as:

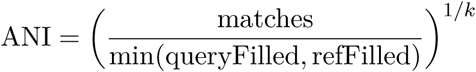

where queryFilled and refFilled are the number of non-empty registers in each sketch. Note that this formula differs from the pairwise WKID formula (Section 5.3): the index provides only match counts, not the lower/equal/higher classification needed for the pairwise denominator. The accuracy characterization in Section 11.2 was measured using pairwise comparison, which is used in practice, and does not directly apply to index-only ANI estimates.

### 6.3 Why This Works for DDL but Not for MinHash, HLL, or a 4-bit Mantissa

DDLIndex exploits two properties of DDL’s encoding:

1. **Discrete value space.** DDL’s 16-bit registers have exactly 65,536 possible values, making the index a finite 2D array. MinHash registers are 64-bit hash values with no natural bucketing — an inverted index would require either an impractically large array (2^64^ cells) or a hash map (losing the strict O(1) lookup property of a direct map, and requiring allocation of memory for keys).
2. **Specificity.** DDL’s structured encoding means that matching values strongly imply shared elements (Section 5.2). Using a pure exponent, or HyperMinHash’s proposed 4-bit mantissa, the index would be dominated by collisions and waste time over brute-force comparison.
3. **Per-bucket independence.** Because each bucket is indexed separately, increasing sketch size (more buckets) does not increase collision rate within the index. Each bucket’s posting list contains exactly *N* entries regardless of the total number of buckets — the value space saturation is determined by *N* and the number of distinct values (65,536), not by *B*. In contrast, MinHash’s flat hash index grows with both *N* and sketch size: doubling the number of hash values per sketch doubles the total keys in the index, increasing collision probability. DDL’s bucketed structure thus scales to larger sketch sizes without degradation.

### 6.4 Index Limitations Compared to BBSketch

DDL’s 16-bit registers with 2,048 buckets effectively constrain approximately 32 bits of the original hash per observation — 10 implied leading zeros (at typical bacterial cardinality), 1 leading one, 10–11 stored mantissa bits, and 11 bucket-selection bits — sufficient to avoid pairwise collisions (0.37 false matches per pair, Section 11.1). However, though random genome pairs have minimal register collisions, individual register values are shared by many sketches across a database of 200,687 references. The index is therefore useful for rapid similarity comparison and for filtering out pairs that share few keys, but cannot support contamination detection.

BBSketch’s contamination analysis counts any hashcode present in query A and some reference C but absent from reference B as a contaminant k-mer with respect to B. This logic requires virtually zero hash collisions — a matching hash must imply an actual shared k-mer, not a coincidence. With 64-bit hash values, BBSketch’s keyspace is large enough to satisfy this assumption. DDL’s 16-bit register values do not: for any register value in query A, there almost certainly exists some sketch C in a 200,687-reference database that shares that value by coincidence — whereas BBSketch’s 64-bit keyspace ensures that such a coincidental match almost certainly does not exist. This makes exclusion-based contamination detection unreliable for DDL.

## 7. Frequency Estimation

### 7.1 Observation Counter

DDL maintains an optional 16-bit counter per register (countArray) that tracks the number of elements achieving the current maximum score:

- When a new element’s score *exceeds* the current register value: counter resets to 1
- When a new element’s score *equals* the current register value: counter increments
- When a new element’s score is *below* the current register value: no change

This counter records the *multiplicity of the winning element* at each register. If the same element is inserted repeatedly and continues to win its bucket, the count grows proportionally to its frequency.

### 7.2 What the Counter Tells You

The counter distribution across buckets characterizes the stream’s frequency profile. This is not a general frequency estimator like Count-Min Sketch — it only tracks the winner at each register, not arbitrary elements. However, for applications where the *distribution* of element frequencies matters (e.g., detecting whether a stream is uniform or skewed), the counter distribution provides a useful signal without maintaining a separate data structure.

### 7.3 Merge Behavior

When merging two DDL sketches, the counter follows the register winner:

- If A’s register exceeds B’s: keep A’s counter and composition
- If B’s register exceeds A’s: keep B’s counter and composition
- If equal: sum the counters (capped at 65,535)

DDL sketches may also be easily shrunk by a factor of 2 (recursively) by treating the upper and lower half of the array as independent sketches and merging them with the same rules, effectively truncating the uppermost bucket bit.

### 7.4 Composition Tracking

DDL optionally maintains a per-register GC content byte (`gcArray`) that records the base composition of the winning k-mer at each register. When a new element’s score exceeds the current maximum, its GC content is stored alongside the new count. This adds 1 byte per register (2 KB at 2,048 buckets). The 5 low-order bits store the precise GC count of the winning k-mer (0–31, sufficient for k ≤ 31; all 8 bits would support k-mers up to length 255). The upper 3 bits are currently unused for k ≤ 31. This could theoretically be employed during comparison to dramatically reduce collision rate, at the cost of increased complexity.

Combined with the frequency counter (Section 7.1), this enables *composition-resolved k-mer frequency histograms* in bounded space. By scanning all registers and binning by count (frequency) and GC content, DDL produces a histogram approximate to what would normally require storing every distinct k-mer and its count — an operation that takes unbounded memory proportional to the number of distinct k-mers. DDL produces a similar histogram shape from a fixed-size sketch.

Figure 2 demonstrates this on a mixed bacterial sample: simulated reads from *Profundibacter amoris* (15× coverage, ∼58% GC) and *Citricoccus* sp. CH26A (40× coverage, ∼72% GC), generated by RandomReadsMG (Bushnell 2025), are processed in a single pass by `loglog.sh`. The resulting k-mer frequency histogram shows two distinct peaks at the expected coverage depths, with GC coloring that visually separates the two organisms. The low-GC peak (blue, ∼58%) at depth 15 corresponds to *Profundibacter*, while the high-GC peak (red/brown, ∼72%) at depth 40 corresponds to *Citricoccus*.

**Figure 2:**
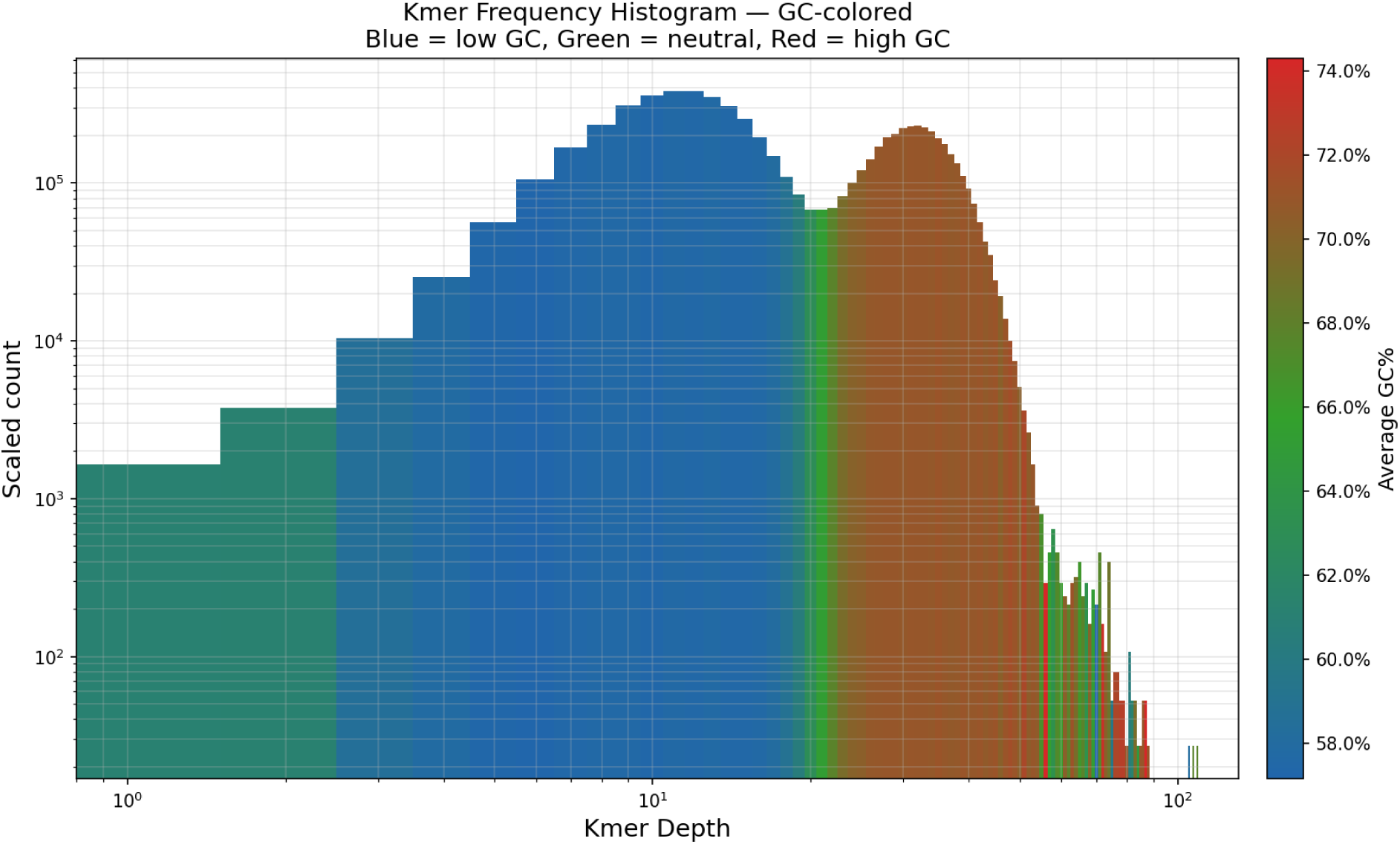
K-mer frequency histogram with GC-content coloring from a mixed bacterial sample. Two species at 15× and 40× coverage are visually separated by GC content (blue = low GC, green = neutral, red = high GC). Generated by `loglog.sh` in a single pass using DDL’s per-register frequency counter and GC tracker, in bounded memory.

This kind of composition-resolved frequency analysis is useful for metagenomics (detecting organisms in mixed samples), quality control (identifying contamination), and coverage analysis (distinguishing GC-biased distributions).

The general principle — frequency distribution with per-element metadata in bounded space — applies beyond bioinformatics to any domain where stream elements carry categorical attributes alongside their identity.

## 8. Parameter Configuration

DDL’s accuracy, sensitivity, and memory depend on four configurable parameters: register width (bits per register), exponent/mantissa split, bucket count (*B*), and k-mer length (*k*). This section discusses the tradeoffs and recommends configurations for different use cases.

### 8.1 Register Width and Encoding

The 16-bit register width was chosen for three reasons:

1. **Hardware alignment.** 16-bit values map to Java `char` (unsigned 16-bit integer), enabling direct array storage without packing overhead, and SIMD-accelerated comparison (16 registers per AVX-256 instruction).
2. **Indexing.** The 65,536 possible values per bucket enable an efficient inverted index (Section 6) — large enough for low false-positive rates, small enough for a direct-mapped array. 6-bit registers (64 values) are too coarse; 64-bit registers (2^64^ values) are too sparse.
3. **Comparison specificity.** 16 bits accommodate a mantissa wide enough for ∼1,000× lower false-match rates than LogLog (Section 11.1), while maintaining sufficient exponent range for cardinality estimation.

DDL uses 2.67× the memory of LL6 (16-bit vs. 6-bit registers, or 4 KB vs. 1.5 KB at 2,048 buckets).

### 8.2 Exponent/Mantissa Split

The exponent/mantissa split is configurable at runtime (`exponent=5` or `exponent=6`; default 6). 6 exponent bits support NLZ values 0–63, covering the full 64-bit hash range without overflow — appropriate for metagenomes and raw read libraries where cardinality is unbounded. 5 exponent bits support NLZ values 0–31, sufficient for individual genomes (cardinalities typically under 10 billion unique k-mers), while freeing one bit for an 11-bit mantissa that halves the per-register false-match probability from ∼1/1,024 to ∼1/2,048 (Section 11.1).

With a 5-bit exponent, elements with NLZ > 31 are masked to 5 bits, wrapping to a low score and typically losing the per-bucket maximum comparison. Since these elements are rare at genomic cardinalities, this has no measurable impact on sketch quality; the bucket is instead won by the next-best element with NLZ ≤ 31. The 5e+11m configuration achieves 96.3% index efficiency on the k=25 RefSeq database (Section 11.7); 6e+10m is slightly lower due its higher collision rate.

### Empirical bit usage

Figure 3 shows the frequency of 1-bits at each position across all 383 million non-empty registers in the RefSeq DDL database (200,687 sketches, 2,048 buckets each).

**Figure 3:**
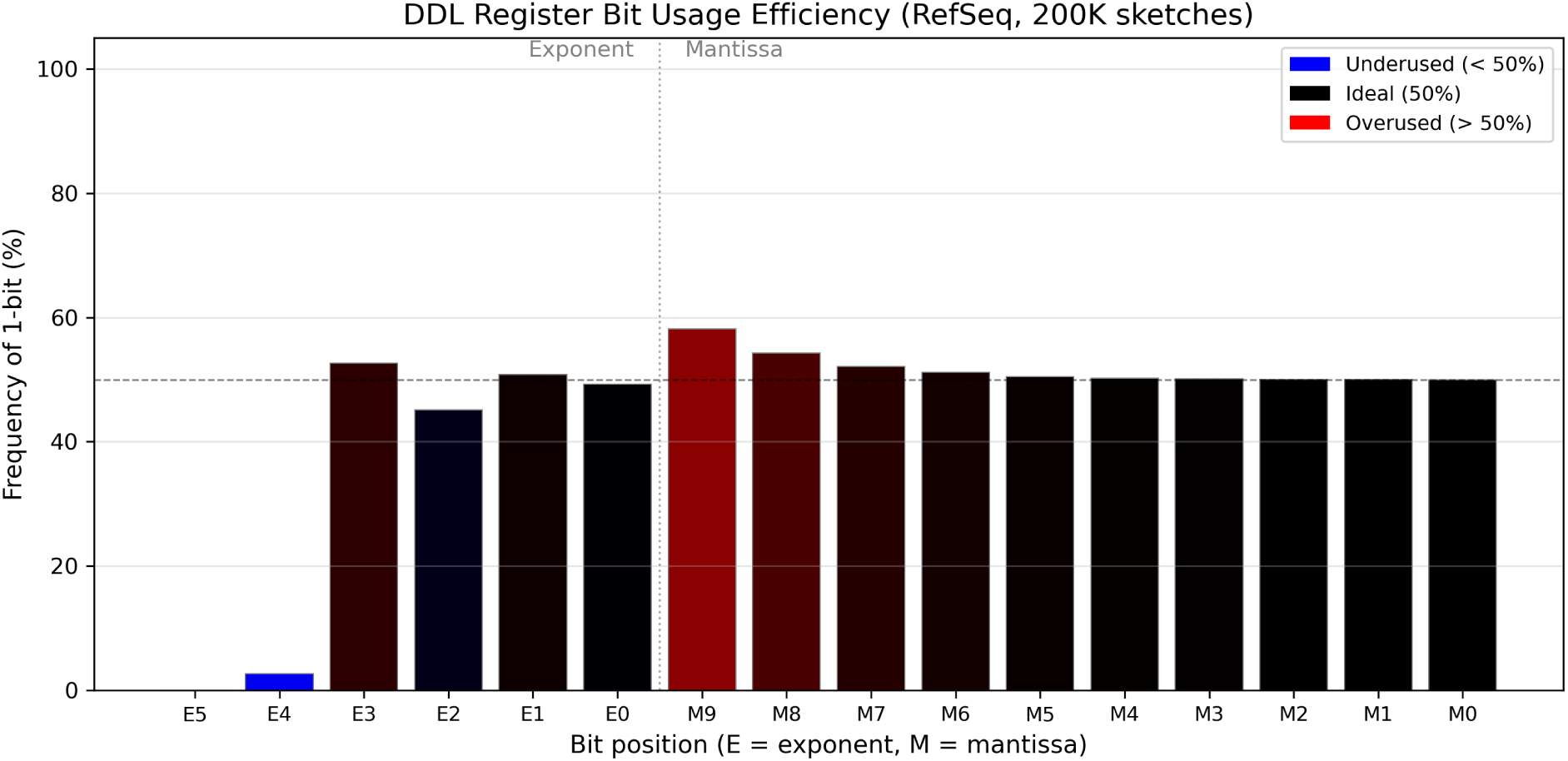
Per-bit 1-frequency across 383M non-empty RefSeq registers. E = exponent (NLZ) bits, M = mantissa bits. Blue indicates underuse (below 50%), red indicates overuse (above 50%), black indicates ideal (50%). The dashed line marks the 50% ideal.

The top exponent bit (E5) is set in only 461 out of 383,446,527 registers (0.0001%), contributing 0.00002 bits of entropy — effectively unused. E4 is set in 2.7% of registers, contributing 0.18 bits. Together, these two bits carry 0.18 bits of information — less than a single ideal bit. The remaining exponent bits (E3–E0) and all mantissa bits (M9–M0) operate near 50%, with total entropy per register of 14.14 bits out of a 16-bit maximum (88.4% efficiency). This confirms that the 5+11 split loses negligible information for genome-sized inputs.

### 8.3 Bucket Count

Increasing the bucket count *B* improves both cardinality accuracy (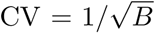; Section 4.1) and comparison sensitivity by lowering the noise floor — the number of false register matches expected between unrelated sketches. At fixed k-mer length, the theoretical expected shared registers between two genomes at a given ANI is *B ×* ANI*^k^*. The noise floor — below which this signal is indistinguishable from random collisions — decreases proportionally with *B*:

The absolute noise floor (matching register count) increases with *B* — 2.09, 2.97, 3.58, 5.01 at *B* = 1,024 / 2,048 / 4,096 / 8,192 — because more buckets create more opportunities for coincidental matches. However, the per-bucket collision rate (noise floor / *B*) decreases: 0.20%, 0.14%, 0.087%, 0.061%, dropping by approximately one-third per doubling of *B*. Since the signal (expected matching registers from genuine similarity) grows linearly with *B*, increasing *B* extends the minimum detectable ANI at the cost of doubled sketch memory.

### Noise floor and reliability floor

We define the *noise floor* as the mean of the top 0.1% of matching register counts between 499,500 unrelated random sketch pairs (Figure 4). The *reliability floor* is double the noise floor, rounded up to the next integer — a conservative threshold above which a match count is overwhelmingly unlikely to arise from random collisions. At *B* = 2,048 with 5e+11m encoding, the noise floor is 2.97, giving a reliability floor of 6 matching registers. The ANI at which the theoretical expected matches (*B* × ANI*^k^*) equal these thresholds under the uniform random substitution model (worst case) is 77% for the noise floor and 79% for the reliability floor at *k* = 25.

**Figure 4:**
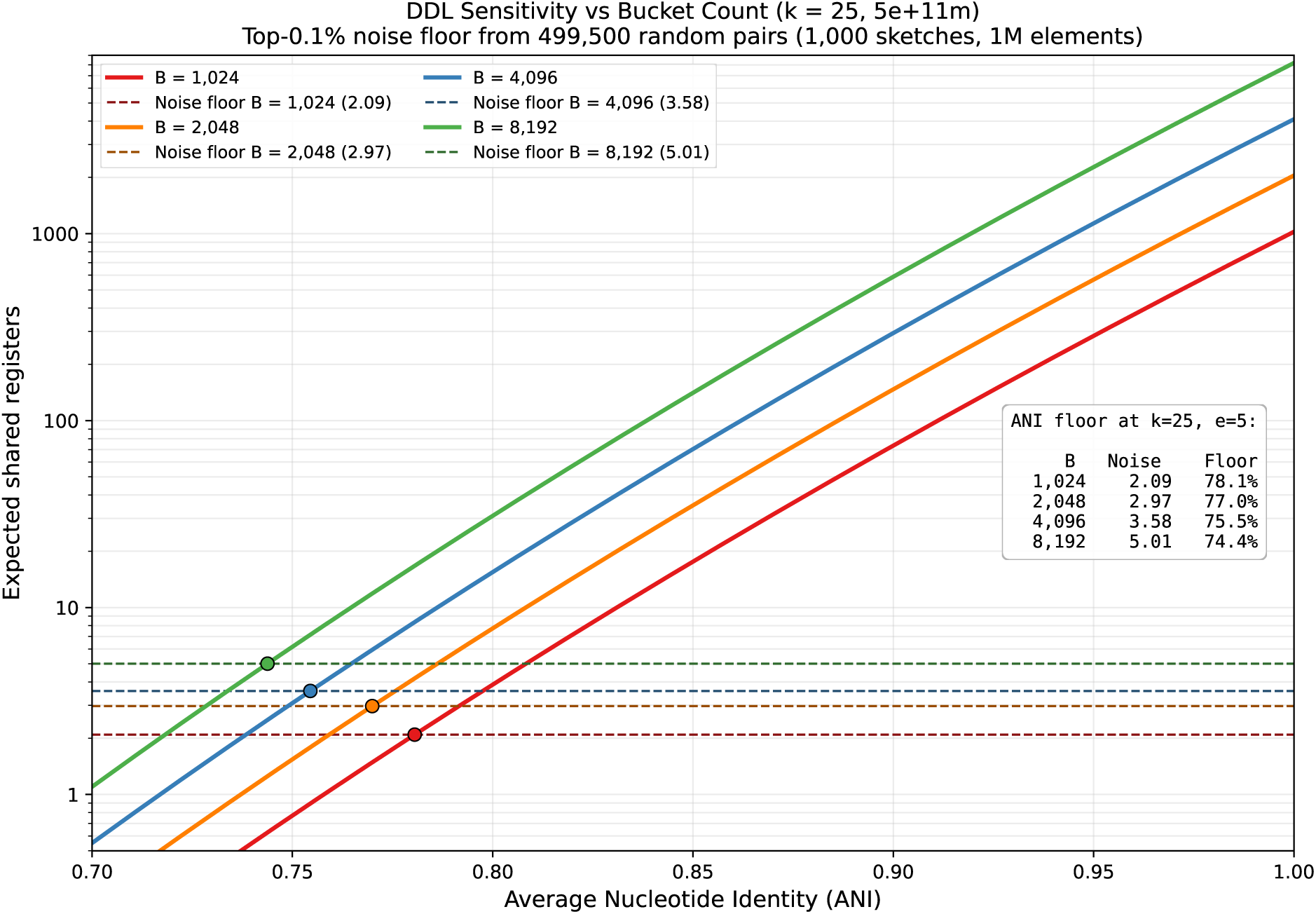
ANI noise floor vs. bucket count at k=25, exponent=5. The noise floor (horizontal lines) is the average of the top 0.1% of matching register counts between 1,000 unrelated random sketches (1M elements each, 499,500 pairs).

The reliability floor of 6 is justified by the false-positive rate: with an empirical per-bucket collision probability of 0.17/2,048 ≈ 8.3 × 10^−5^ (from the 5e+11m collision benchmark, Section 11.1), the probability of ≥ 6 coincidental matches in 2,048 independent buckets is *P* (Binomial(2048, 8.3 × 10^−5^) ≥ 6) ≈ 2.9 × 10^−8^ — approximately 1 in 35 million per unrelated same-size pair. Against a database of 200,687 variable-size references, the expected number of false positives per query at the 6-match threshold is below 0.006.

### 8.4 K-mer Length

Shorter k-mers increase sensitivity at low ANI because more k-mers survive mutation: at 80% ANI, the k-mer survival probability is 0.80^31^ ≈ 0.001 at k=31 versus 0.80^25^ ≈ 0.004 at k=25 — a 4× increase in expected matching registers. Yet shorter k-mers also increase the probability of both coincidental and ultra-conserved matches between unrelated and distantly related genomes, raising the noise floor for ANI calculation. The coincidental match rate is easy to calculate assuming random sequences, and remains low at k=25. However, real sequences are not random; the expected rate of shared long k-mers between distantly-related organisms, as a factor of k, is impossible to derive theoretically and varies by clade. Thus, reducing k has a predictable effect of increasing sensitivity but an unpredictable effect on increasing noise, and the optimal value is subjective: “as short as possible while retaining sufficient specificity for the purpose”.

### 8.5 Recommended Configurations

The default configuration — 6-bit exponent, 10-bit mantissa, B=2,048, k=31 — is designed for maximum generality. The 6-bit exponent handles unbounded cardinality (metagenomes, raw read libraries, filesharing streams), the 2,048-bucket sketch fits in 4 KB, and k=31 provides high specificity for closely related genomes.

For genome-to-genome comparison against a bounded reference database (e.g., RefSeq), the configuration exponent=5, mantissa=11, B=2,048, k=25 provides improved sensitivity and specificity: the 11-bit mantissa halves the false-match rate, and k=25 extends the detectable ANI range to approximately 79%. This is DDLCompare’s default configuration. A denser B=4,096 variant further reduces the noise floor for applications requiring sensitivity at deeper phylogenetic distances, at the cost of doubled sketch size.

### 8.6 Index Size Optimization

The inverted index (Section 6) dominates DDL’s memory footprint in indexed mode — the index requires 9.7 bytes per key compared to 2.4 bytes per key for the sketches alone (Section 11.7). Two orthogonal optimizations can reduce this:

### Partial bucket indexing

Instead of indexing all *B* = 2,048 buckets, the index can be built from only the first *B’*<*B* buckets, reducing index memory and query scan time from O(*B*) to a constant, at the cost of proportionally fewer match opportunities per query. For organisms with high WKID, using *B*/2 = 1,024 buckets provides sufficient signal for identification in half the memory; for distant organisms near the detection threshold, sensitivity is reduced but not eliminated.

### 15-bit key masking

The bit frequency analysis (Section 8.2) shows that bit 15 (E5) is set in only 461 out of 383 million registers when using a 6-bit exponent — effectively zero information. Masking register values to 15 bits before indexing reduces the value space from 65,536 to 32,768 entries per bucket, doubling the index density by eliminating null pointers. The 461 affected keys are folded into their 15-bit neighbors, causing a negligible increase in false matches. Alternatively, registers with bit 15 set can simply be excluded from the index.

### Combined effect

An index with 1,024 buckets and 15-bit keys (`int[1024][32768][]`) uses less than half of the memory of the full index (`int[2048][65536][]`) and processes queries in half the time while retaining full precision of sketch comparisons. This tradeoff is attractive for memory-constrained deployments or when query throughput is prioritized over marginal sensitivity at low ANI.

## 9. Taxonomic K-mer Blacklisting

DDL’s inverted index (Section 6) eliminates 96.3% of brute-force comparisons by looking up each query register in a (bucket, value) → sketch ID map. However, taxonomically conserved k-mers — rRNA operons, housekeeping genes, transposable elements — can win bucket competitions across phylogenetically distant organisms, producing identical register values in thousands of unrelated sketches. These “hot” register values generate dense posting lists in the index, inflating the number of candidate pairs that must be evaluated by full pairwise comparison.

Taxonomic k-mer blacklisting addresses this by identifying and excluding the most broadly shared k-mers from sketch construction. The filtering is integrated into `hashAndStore()` (Section 3.2): after the early exit mask (Section 3.2) and the `score <= register[bucket]` check reject the vast majority of elements, a single hash probe against a static `LongHashSet` of blacklisted k-mers rejects conserved k-mers before they can update a register. The check applies only to the rare elements that survive both prior filters, making the per-element cost negligible. Pre-built reference databases have blacklisting baked into the sketches; query sketches are filtered at construction time by autoloading the appropriate blacklist from the `resources/` directory.

### 9.1 Blacklist Generation

To identify which k-mers to blacklist, DDL sketches are built with literal k-mer storage (`kmers=t`), which records the canonical packed 2-bit k-mer that won each bucket in a parallel `long[]` array alongside the register values. This adds 16 KB per sketch (2,048 × 8 bytes) during blacklist generation only; production sketches omit k-mer storage.

`DDLBlacklistMaker` loads a set of DDL sketches with stored k-mers, promotes each sketch’s NCBI TaxID to genus level via the taxonomy tree, and counts the number of distinct genera containing each k-mer across the entire database. Genus-level promotion prevents strain-level redundancy from inflating counts — 50 strains of *E. coli* count as one genus, not 50. K-mers exceeding a threshold at any taxonomic level are flagged for blacklisting. Thresholds are applied independently at five levels — genus, family, order, class, phylum — because each level captures a different scale of uninformativeness: a k-mer shared by 50 families may indicate clade-specific signal (e.g., highly conserved ribosomal sequence), while a k-mer shared by 50 phyla is not useful for low-level taxonomic classification. The per-level blacklists are merged with deduplication (`MergeDDLBlacklists`), producing a single FASTA file of blacklisted k-mer sequences.

The generation process uses two iterations. In the first pass, oversize sketches are built without a blacklist, and a preliminary blacklist is generated from the most broadly shared k-mers. In the second pass, sketches are rebuilt with the preliminary blacklist active, and a refined blacklist is generated from the cleaned data — this catches k-mers that were masked in the first pass by higher-scoring k-mers occupying the same buckets. The second pass adds approximately 10% additional k-mers; a third pass adds negligibly more.

To the authors’ knowledge, taxonomic k-mer blacklisting at the sketch level — filtering k-mers by their taxonomic breadth before sketch construction, rather than post-hoc during comparison — has not been described for any MinHash or LogLog-family sketch. BBSketch (Bushnell 2016) maintains a similar blacklist of taxonomically uninformative hash values identified by their prevalence across reference sketches, operating in full 64-bit hash space. DDL’s blacklist operates on literal k-mers (canonical 2-bit-packed sequences), enabling the same blacklist to be applied across sketches built with different hash seeds or parameters.

### 9.2 Ribosomal Database Blacklists

For SSU/ITS ribosomal databases (k=19, 128 buckets, exponent=4), per-type blacklists were generated from the constituent databases and merged:

**Table 3:** Ribosomal blacklist composition. Only 7 blacklisted k-mers are shared across all three gene types, reflecting distinct conserved cores.

| Database | Sequences | Thresholds (fam/ord/cls/phy) | K-mers |
| --- | --- | --- | --- |
| 16S (prokaryotic SSU) | 221,462 | 160 / 80 / 32 / 18 | 1,219 |
| 18S (eukaryotic SSU) | 55,310 | 190 / 70 / 25 / 12 | 2,758 |
| ITS (fungal) | 35,429 | 200 / 70 / 24 / 8 | 217 |
| **Combined** | **312,201** | — | **4,230** |

The noise floor (average shared keys at class+ taxonomic distance) was reduced 10–29× depending on gene type: ITS from 7.18 to 0.25 (29×), 18S from 10.92 to 0.45 (24×), 16S from 10.85 to 1.06 (10×). All-to-all indexed comparison of 312,201 sketches at 32 threads was 2.6× faster with the blacklist (114.5s → 44.0s), while the number of real alignments was virtually unchanged (6,089,276 vs. 6,088,203) — the blacklist overwhelmingly eliminated spurious index candidates, not genuine matches.

### 9.3 Genome Database Blacklists

For the RefSeq genome database (k=31, 2,048 buckets, exponent=5), blacklists were generated from 200,687 sketches across all 12 RefSeq clades (archaea, bacteria, fungi, invertebrate, mitochondrion, plant, plasmid, plastid, protozoa, vertebrate_mammalian, vertebrate_other, viral). Genus-promoted counting with thresholds of 250 (genus), 50 (family), 20 (order), 10 (class), and 5 (phylum) distinct genera per k-mer produced a combined blacklist of 23,020 k-mers after two-pass iterative refinement and deduplication.

Without blacklisting, the class average unexpectedly exceeds the order average in both the unmerged (32.12 vs. 19.68) and merged (15.49 vs. 11.30) databases — an inversion of the expected monotonic decrease with taxonomic distance. With the 23,020 k-mer blacklist, the inversion is eliminated and the expected ordering is restored (e.g., unmerged: order 6.17 > class 2.33 > phylum 0.23); the noise floor drops 13× (11.51 → 0.86 unmerged, 5.98 → 0.47 merged).

Genus-level signal is preserved: average shared keys between congeneric pairs decrease by only 1.5% (162.73 → 160.37 unmerged, 182.65 → 181.39 merged), confirming that the blacklisted k-mers contribute almost exclusively to noise rather than signal.

### Index performance

All-to-all indexed comparison of 200,687 unmerged RefSeq sketches at 32 threads completed in 68.9s with the blacklist vs. 103.4s without (1.50× speedup). At 128 threads, the speedup was 1.36× (28.0s vs. 38.1s) — smaller because the higher thread count shifts the bottleneck from compute to memory bandwidth, partially masking the reduced comparison count. Index efficiency (fraction of brute-force comparisons eliminated) improved from 95.3% to 96.7%, corresponding to a 30% reduction in the absolute number of indexed comparisons (1.91 billion → 1.34 billion).

The merged database showed similar speedups: 1.47× at 32 threads (67.0s → 45.6s) and 1.31× at 128 threads (26.9s → 20.5s).

### Blacklisted k-mer entropy

Blacklisted k-mers are not low-complexity repeats. The mean Shannon entropy (computed over sub-3-mers, normalized to [0, 1]) of the 23,020 blacklisted k-mers is 0.9195, compared to 0.8903 for all k-mers appearing in ≥ 5 genera — blacklisted k-mers are *higher* entropy than the background. 95% of blacklisted k-mers have entropy ≥ 0.85; only 26 fall below 0.70 and only 10 below 0.50. The blacklisted population consists predominantly of conserved functional genes (rRNA operons, tRNA synthetases, ribosomal proteins, housekeeping enzymes) whose sequence conservation across deep phylogenetic distances makes them taxonomically uninformative despite their high sequence complexity. This distinguishes DDL blacklisting from low-complexity filtering (e.g., DUST or entropy masking), which targets repetitive sequence; the two mechanisms are complementary.

#### 9.3.1 Genome-Optimized Database (k=25, B=2,048)

A genome-optimized RefSeq database at k=25, B=2,048, exponent=5 is available as of v39.91 and is the default configuration for DDLCompare. At 80% ANI, the k-mer survival probability is 0.80^25^ ≈ 0.004, achieving a reliability floor (Section 8.3) of approximately 79% (k=25, 5e+11m) at B=2,048. A denser B=4,096 variant is also available for applications requiring lower noise floors.

The tradeoff between k-mer length and ANI resolution is shown in Figure 5: at B=2,048, the theoretical expected shared registers (B × ANI^k) drops below the empirical noise floor at progressively higher ANI as k increases — approximately 71% for k=19, 73% for k=21, 77% for k=25, and 81% for k=31 (5e+11m encoding, noise floor 2.97). Below this floor, genuine matches become difficult to distinguish from random collisions. At fixed k=25, increasing B shifts the noise floor downward (Figure 4, Section 8.3), with the per-bucket collision rate dropping by approximately one-third per doubling of *B*.

**Figure 5:**
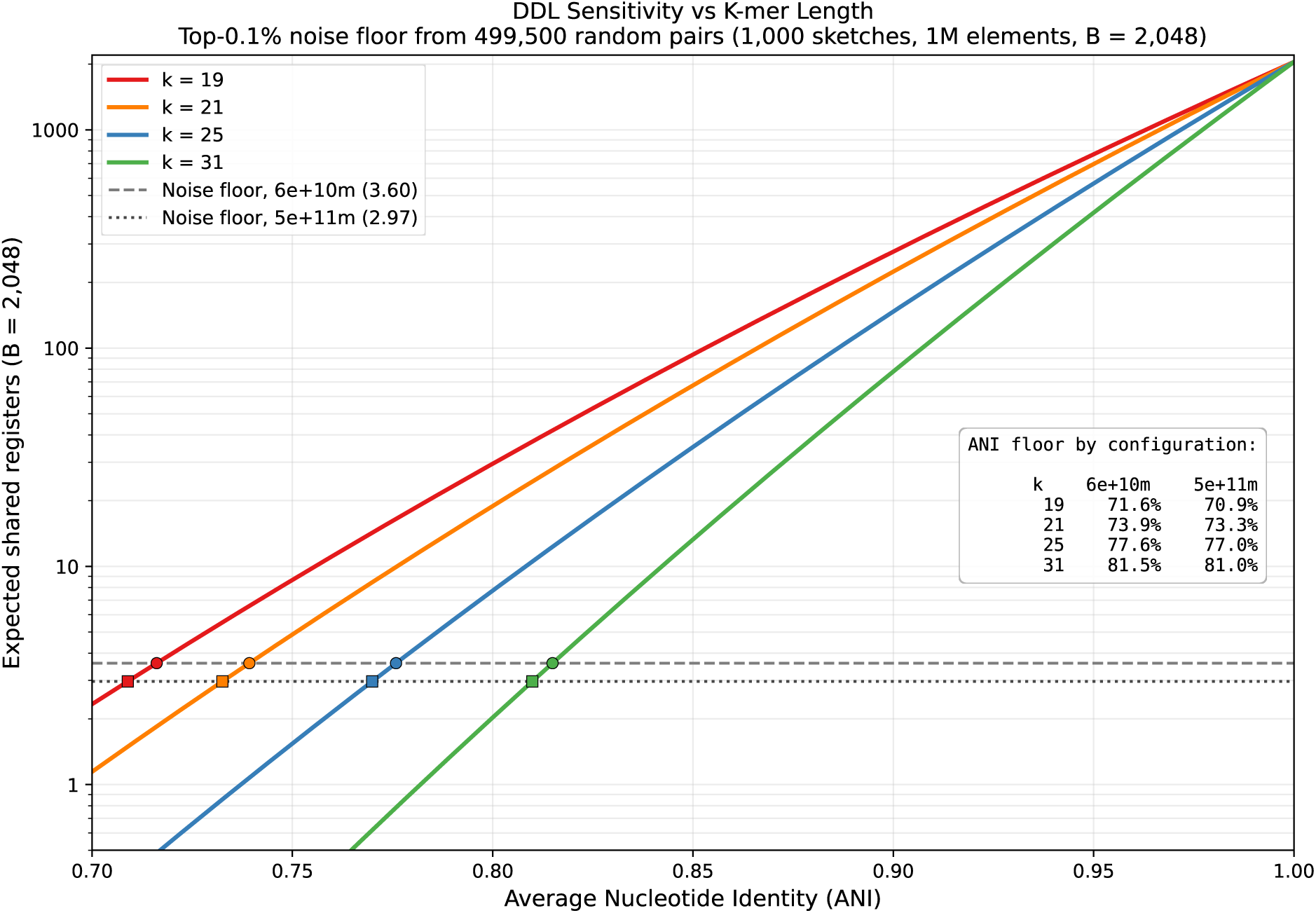
Expected shared registers vs. ANI for k=19, 21, 25, and 31 at B=2,048, with empirical noise floors (horizontal lines). The noise floor is the average of the top 0.1% of matching register counts between 1,000 unrelated random sketches (1M elements each, 499,500 pairs). At the 5e+11m encoding (noise floor 2.97), the theoretical curve crosses the noise floor at approximately 81% for k=31, 77% for k=25, 73% for k=21, and 71% for k=19.

The k=25 blacklist contains 1,651 k-mers (genus ≥ 1,600, family ≥ 270, order ≥ 110, class ≥ 40). Shorter k-mers are less likely to be conserved identically across deep phylogenetic distances, so fewer exceed the taxonomic breadth thresholds. The blacklisted k-mers are high-entropy conserved genes, not low-complexity sequence.

The class-order inversion (class > order in shared keys) persists in the unmerged blacklisted database (20.89 vs. 17.28). This inversion is primarily a database artifact: organelle and plasmid sketches share their host’s TaxID, inflating class-level counts via duplicate signal, and incomplete taxonomic annotations leave some records without intermediate ranks (e.g., missing order assignments), biasing the class average upward. The inversion nearly resolves in the merged database (9.88 vs. 9.55), where absorbing organelle and plasmid sketches into their host genomes removes the duplicate signal. In all configurations, both class and order averages fall well below any meaningful ANI threshold, so the inversion does not affect taxonomic assignment.

The noise floor decreases from 2.61 (no blacklist) to 1.21 (blacklisted unmerged) to 0.62 (blacklisted merged) — a 4.2× total reduction. Species-level signal is preserved: average shared keys change by <0.02% (1,251.56 → 1,251.42).

### 9.4 Blacklist as Infrastructure

The blacklist check is integrated into `DynamicDemiLog.hashAndStore()` and applies to all tools that construct DDL sketches — DDLCompare, DDLWriter, QuickClade, and the ribosomal tools (FindSSU, SSUCom-pare, SSUServer). Each tool autoloads its database-appropriate blacklist from the `resources/` directory at startup: `genomeDDLBlacklist_k25e5b4096.fa.gz` (1,651 k-mers, 40 KB) for the k=25 genome database, and `riboDDLBlacklist.fa.gz` (4,230 k-mers, 27 KB) for ribosomal databases. Pre-built reference sketches have blacklisting baked in; the autoloaded blacklist filters only query sketch construction, ensuring consistency between query and reference. The blacklists are distributed with BBTools, so local queries to remote databases using QuickClade or FindSSU will generate sketches with blacklists applied.

## 10. Methods

### 10.1 Collision Benchmark (Synthetic)

The collision test uses purely synthetic random input to evaluate per-register specificity independent of any application domain. DDLBenchmark generates random 64-bit values via Xoshiro256 PRNG (`FastRandomXoshiro`) and inserts them into CardinalityTracker instances. All comparisons use the `bucketValues()` interface.

100 independent sketches are built from random streams of 4 million elements each (representative of an average bacterial genome). All-pairs comparison (4,950 pairs) counts the number of registers that match (equal stored value) despite the sketches being constructed from independent random streams. Any match is a false positive.

### 10.2 ANI Benchmark (Nucleotide Sequences)

The ANI accuracy test uses real nucleotide sequences with controlled mutation. A 1 MB random genome (GC-neutral) is generated with `randomgenome.sh`, then mutated copies are created using `mutate.sh` (BBTools) at 121 per-nucleotide substitution rates from 0.1% to 30.0% in 0.25% steps (design ANI = 99.9% to 70.0%). Substitutions are uniform random across the four bases; 99% of mutations are substitutions and 1% are indels.

For each mutation rate, three ANI measurements are taken:

1. **Measured ANI** — alignment-based identity computed by quantumaligner.sh (BBTools), which performs exact dynamic programming alignment and reports the fraction of matching nucleotides; measured ANI averages higher than design ANI due to coincidental alignment matches of historically unrelated positions and plateaus at ∼54% (Bushnell 2026c).
2. **DDL ANI** — DDL sketches are built from the reference and mutant genomes by hashing all *k*-mers via the standard `hash(Read)` pipeline. WKID and ANI are estimated from bucket comparison as described in Section 5.3.
3. **LL6 ANI** — identical pipeline using 6-bit LogLog (LL6) sketches instead of DDL, with the same comparison and ANI estimation.

This three-way comparison isolates the effect of the mantissa on a biologically realistic mutation model: design ANI is the engineered per-nucleotide identity, measured ANI is the alignment truth, and sketch-based ANI estimates are what the user would compute in practice.

### 10.3 Speed and Scaling Benchmarks (Synthetic)

#### Speed mode

50 replicate sketches are built sequentially on a single thread, each from 5 million random hash elements, with 3 warmup iterations discarded. Wall-clock throughput is measured in adds per second.

#### Scaling mode

A reference sketch from 5 million random elements is compared against query sketches of sizes 2,000 to 64,000,000 from independent random streams, measuring false matches and spurious WKID.

### 10.4 Sketch Parameters

All sketches use *B* = 2,048 buckets. Synthetic benchmarks (Sections 10.1, 10.3, 10.5) use *k* = 31 and the default 6e+10m encoding, since they operate on random hash values where k-mer length does not affect results. The ANI accuracy benchmark (Section 10.2) uses *k* = 25 and exponent=5, matching the genome-optimized default. DDL uses 16-bit registers (NLZ exponent + inverted mantissa). LL6 uses standard 8-bit registers storing only the 6-bit NLZ. Both use the same hash function: Thomas Wang’s 64-bit integer hash (`hash64shift`) (Wang 1997), applied after XOR with a configurable seed. For k-mer inputs, k-mers are first encoded as canonical 2-bit-encoded 64-bit integers, then hashed. SetSketch uses the same hash function and bucket count with base *b* = 1.001 and 16-bit registers.

### 10.5 Cardinality Accuracy Benchmark

DDL and LL6 cardinality estimation accuracy was compared using DDLCalibrationDriver2 (BBTools), which creates many independent estimator instances and feeds each a stream of random 64-bit values generated by Xoshiro-256, measuring fractional estimation error at logarithmically-spaced cardinality thresholds from 1 to *B* × maxMult.

Three calibration runs were performed:

1. **DDL at 2,048 buckets (4 KB)** — tests DDL’s MeanM mantissa-based estimators
2. **LL6 at 2,048 buckets (2 KB)** — same bucket count, tests whether the mantissa improves per-register accuracy
3. **LL6 at 4,096 buckets (4 KB)** — same memory footprint, tests whether DDL’s extra information per register is worth more than LL6’s extra buckets

From these runs, four estimators are compared: the production Hybrid estimator for DDL and LL6 (which blends Linear Counting with HLL or MeanM), and the raw HLL estimator for LL6 at both bucket counts (which shows the unblended LC-to-HLL transition behavior).

Each configuration used 524,288 independent estimator instances, maxMult = 8,192 (maximum cardinality = *B* × 8,192), and correction factor tables enabled (cf=t). Hardware details are in Section 10.9. Error metrics reported include log-weighted, width-weighted, and count-weighted mean absolute error, peak absolute error, and mean signed error across all cardinality thresholds.

### 10.6 Database Query Benchmark

Three tools were benchmarked for database query performance: DDLCompare (DDL-based), CompareSketch (BBTools’ MinHash implementation), and Mash v2.3 (Ondov et al. 2016). Each tool searched its native reference database: DDLCompare against 200,687 DDL sketches (411M keys, 595 MB compressed on disk), CompareSketch against 85,155 MinHash sketches (1.21B keys, 9.8 GB), and Mash against 91,282 MinHash sketches (91.3M keys, 720 MB). The query for all tools was 500,000 metagenomic read pairs (151 Mbp decompressed FASTQ). Sketch construction speed was measured separately on 14.7 million reads (2,623 Mbp) from a bacterial isolate to ensure timing was dominated by hashing rather than JVM startup.

Each benchmark was decomposed into phases: query sketch construction (Mbp/s), reference loading (MKeys/s), brute force comparison (MKeys/s and queries/s), and — where supported — index construction and indexed query time. All input files were decompressed to local storage before benchmarking to eliminate I/O variability. Timings were measured at both 1 and 32 threads to characterize scaling behavior.

Memory was measured by heap bisection: finding the minimum Java heap (`-Xmx`) at which the program runs without crashing. Mash uses memory-mapped binary I/O; its memory reflects file size per key, not Java heap.

For brute force comparison, 1,000 pre-extracted queries were compared against the full reference database. DDLCompare used multi-query mode (`queryfile=`); CompareSketch used `in=` with pre-sketched queries; Mash used pre-sketched queries via `mash dist`.

### 10.7 Completeness Benchmark

The *E. coli* K-12 genome (4.64 Mbp) was truncated to 5%, 10%,…, 100% of its length, yielding 20 size fractions. Each fraction was sketched as a DDL (k=25, B=2,048) and compared against the full-length reference sketch. To test robustness under mutation, the experiment was repeated at four ANI levels: 100% (identical sequence), 99% (1% substitution via `mutate.sh`), 95% (5% substitution), and 90% (10% substitution). Completeness was estimated using the formula in Section 5.3.

### 10.8 Genome Size Benchmark

Genome size estimation was tested on a real sequencing library of *Micromonospora rosaria* (JGI library ABXYZ_GAGATCTATG-AGGTCGCGCG, known genome size 7.28 Mbp, 178 MB compressed FASTQ, approximately 17× coverage). DDL (`loglog.sh`) was compared against KmerCountExact (`kmercountexact.sh`, exact hash table) and BBNorm (`khist.sh`, Count-Min Sketch with 1 GB heap). All three tools produce k-mer frequency histograms from which genome size is estimated by a peak-finding algorithm that separates error k-mers from genomic k-mers.

DDL’s bucket-count scaling was additionally tested at 2,048 to 524,288 buckets (4 KB to 1 MB sketch size) to characterize the relationship between sketch size and estimation accuracy.

### 10.9 Environment

Benchmarks in Sections 10.1–10.5 are single-threaded — each mode runs sequentially on a single core. The database query benchmark (Section 10.6) is measured at both 1 and 32 threads. Speed and database query benchmarks (Sections 10.3, 10.6) ran on an AMD EPYC 7502P (32 cores, 2.5 GHz); calibration benchmarks (Section 10.5) and the 128-thread all-to-all comparison (Section 11.7) ran on single-socket AMD EPYC 9555P nodes (64 cores, 128 threads, 3.2 GHz) via batch submission. All cluster benchmarks used OpenJDK 25 with `jdk.incubator.vector` enabled and 64 GB heap. BBTools version 39.91. Local validation ran on a consumer laptop.

## 11. Experimental Results

### 11.1 Collision Rate

The collision test measures how often registers from unrelated random sketches produce matching stored values — a direct test of per-register specificity.

DDL (6e+10m) achieves a **930× reduction** in false register matches compared to LL6 (Figure 6). SetSketch’s fine-grained logarithmic encoding (*b* = 1.001) eliminates LL6’s dominant-tier problem but still produces 40% more false matches than DDL (6e+10m), because the logarithmic values cluster where hash magnitudes are dense. DDL’s separate mantissa provides effectively random discrimination within each exponent tier — two unrelated registers in the same NLZ tier have only a ∼1/2^*m* chance of matching.

**Figure 6:**
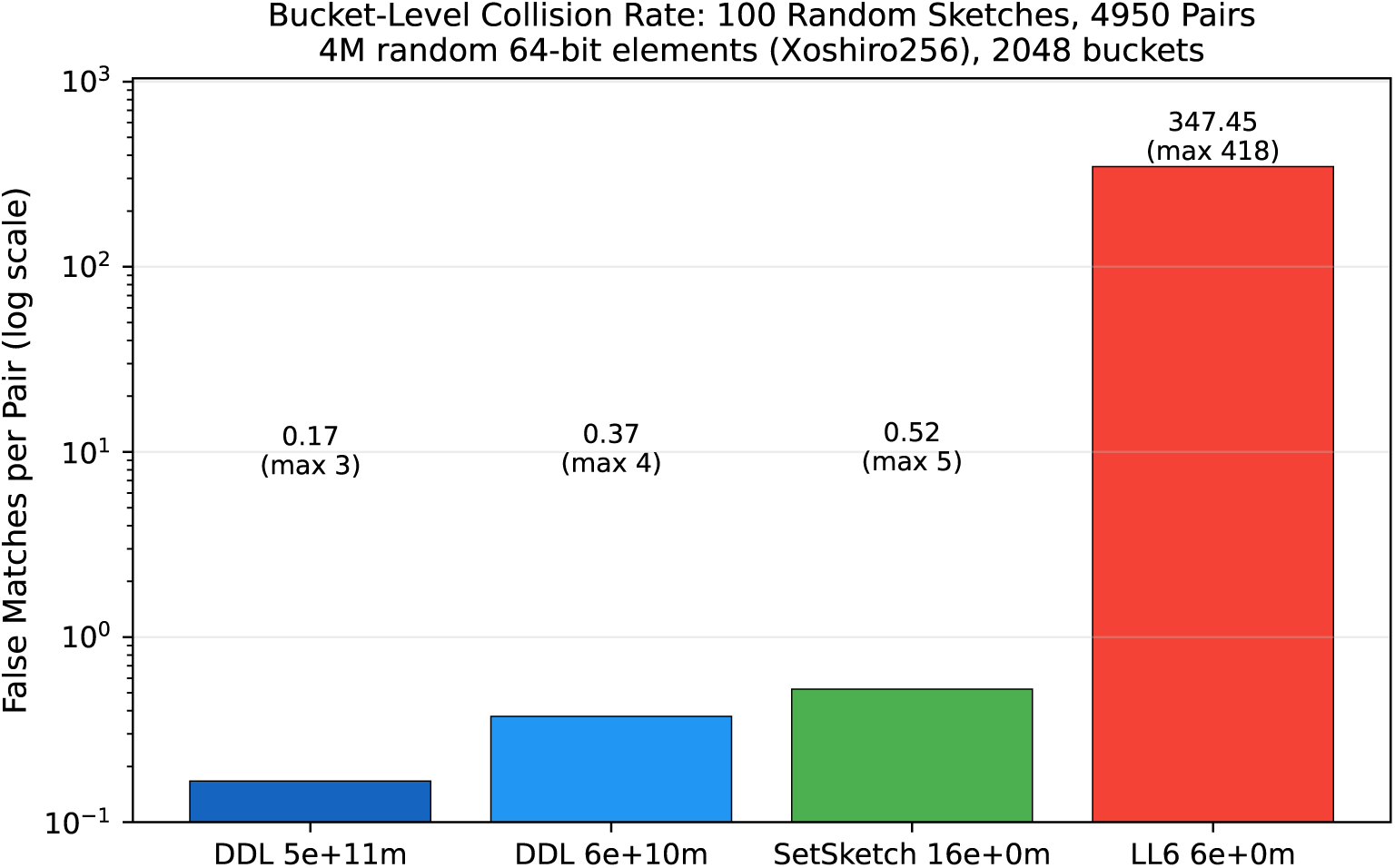
Collision rate comparison: DDL 5e+11m, DDL 6e+10m (default), SetSketch (16-bit exponent, b=1.001), and LL6 (6-bit exponent only). Log scale. 100 random sketches, 4,950 pairs, 4M random elements per sketch (Xoshiro256 PRNG), 2,048 buckets, all 16-bit registers.

The expected false match count can be derived empirically: LL6 averages 347.5 same-register matches per pair (the same-tier collision rate), and the default 10-bit mantissa subdivides each tier into 1,024 values, giving an expected 347.5 / 1,024 ≈ 0.34 false matches — consistent with the observed 0.37.

The 5e+11m configuration (Section 8.2) halves false matches to 0.17 per pair by doubling the mantissa to 2,048 intra-tier values, with no measurable impact on cardinality accuracy. The 6e+10m split remains the default for maximum generality, but 5e+11m is preferable for bounded-size databases such as RefSeq.

### 11.2 ANI Estimation Accuracy

Figure 7 shows ANI estimation accuracy across 121 mutation rates using real nucleotide sequences. Three estimates are plotted against design ANI (the engineered per-nucleotide identity):

- **Measured ANI** (black): alignment-based identity from quantumaligner, representing ground truth. Measured ANI slightly exceeds design ANI due to coincidental alignment matches of unrelated bases.
- **DDL ANI** (blue): WKID^1^*^/k^* from DDL register comparison. DDL tracks measured ANI closely down to approximately 79% design ANI (≥6 matching registers out of 2,048). Below approximately 80%, too few k-mers survive mutation for reliable estimation — at 77% ANI and *k* = 25, the expected k-mer survival probability is 0.77^25 ≈ 0.0009, yielding approximately 2 predicted matching registers.
- **LL6 ANI** (red): same computation from LL6 register comparison. LL6 reports a floor of ∼95% ANI regardless of true identity, because its integer-NLZ registers produce false matches at a rate that dominates the signal. LL6 cannot distinguish related from unrelated genomes.

#### Practical ANI floor

DDL’s ANI resolution at *B* = 2,048 and *k* = 25 is limited by the number of surviving k-mers. The summary table (50 trials per level) shows that DDL tracks measured ANI within 0.1% down to 87% (32.0 matches, *σ* = 0.57%) and remains accurate at 85% (16.8 matches, *σ* = 0.78%). Between 85% and 80%, the standard deviation explodes from 0.78% to 11.3% as average matches drop from 16.8 to 4.2 — a sharp cliff in reliability rather than a gradual decline.

Increasing *B* extends the range: at *B* = 4,096, the lower per-bucket collision rate (0.087% vs. 0.14%) and doubled bucket count yield approximately 2× more matching registers at each ANI level. The theoretical noise-floor crossing (Section 8.3) projects a floor of approximately 73% at *B* = 4,096 and 69% at *B* = 8,192, though these have not been empirically validated.

Note that even a small number of matches (8 or more out of 2,048) provides a strong signal for taxonomic assignment, though not for precise ANI estimation. For comparison, BBSketch uses a floor of 3 matching keys out of sketches averaging 14,220 keys — allowing sensitivity to ∼75% ANI with dual 32- and 24-mers. BBSketch’s sensitivity is further increased by using oversized query sketches to compensate for error k-mers in raw sequence that use sketch space relative to the smaller reference sketches. Oversized queries are possible to implement but more complex with DDL.

For GC-neutral random sequences with no evolutionary relationship, measured ANI converges to approximately 54% — an empirically observed floor that reflects coincidental nucleotide matches in aligned regions.

**Figure 7:**
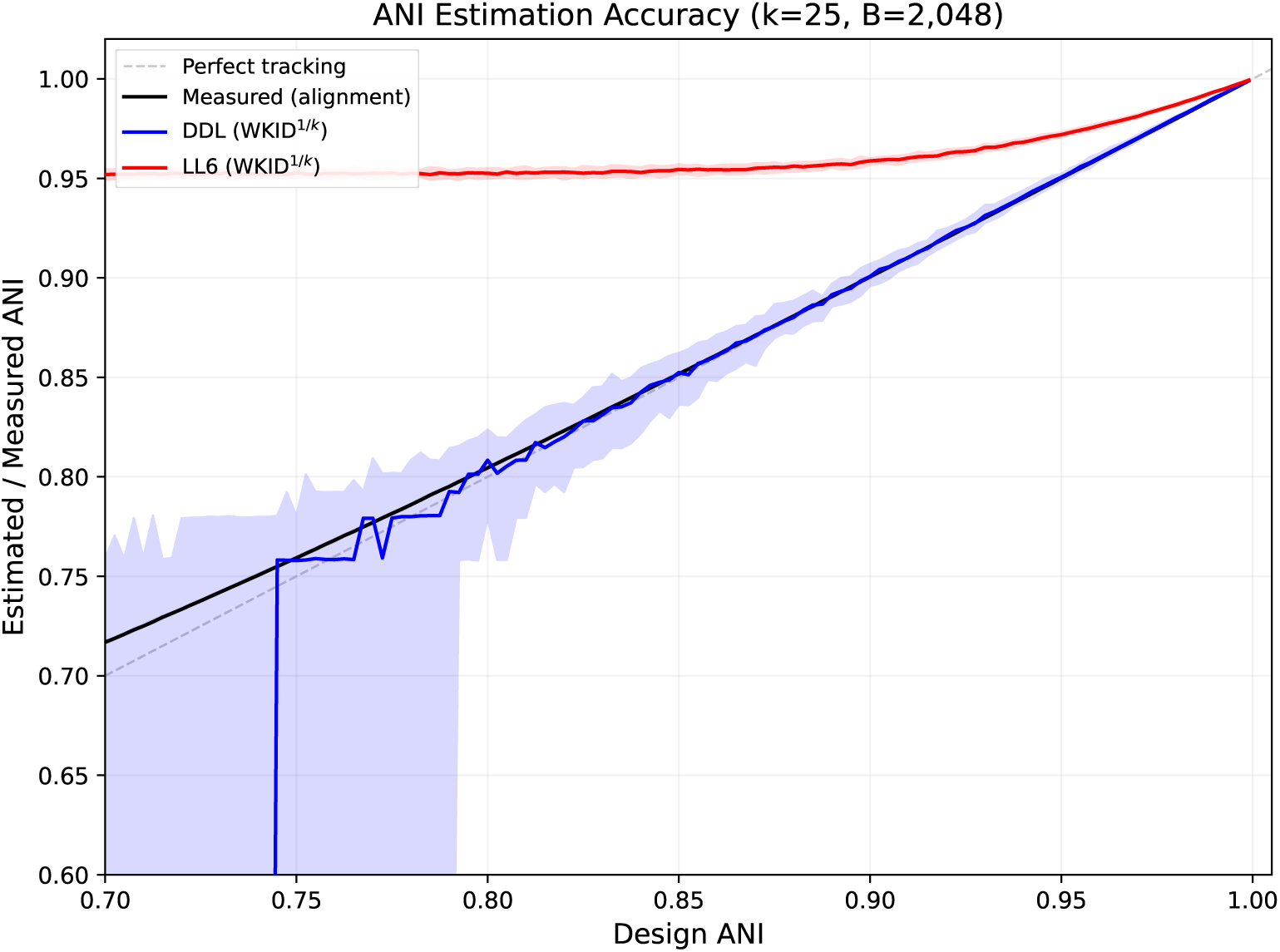
ANI estimation accuracy: design ANI vs measured (black), DDL estimate (blue), and LL6 estimate (red). Scatter points are individual measurements; lines are smoothed curves. 121 ANI levels from 70% to 99.9%, 50 trials each. 1 MB random genome, mutate.sh, 2,048 buckets, k=25, exponent=5.

#### ANI robustness to genome size differences

ANI estimation accuracy depends not only on mutation rate but also on relative genome completeness. Tools that estimate ANI from Jaccard similarity (intersection over union) inherently underestimate ANI when the query and reference differ in size, because the union grows faster than the intersection. Tools that use WKID (matches over the minimum of filled positions) are invariant to size differences — a partial genome produces the same ANI estimate as the full genome, provided sufficient matching registers exist.

Figure 8 demonstrates this by comparing a 99% ANI mutant of *E. coli* K-12, truncated to 1–100% of its original length, against the full reference. DDL and BBSketch both use WKID-based ANI and produce stable estimates (∼99%) across the full completeness range. Mash uses Jaccard-based ANI and underestimates severely at low completeness — reporting 85.5% ANI at 1% completeness for a genome that is actually 99% identical. Even at 50% completeness, Mash underestimates by 2 percentage points. The ANI formula is symmetric, so swapping query and reference produces the same bias. This property is critical for applications such as metagenomic binning, where query completeness is often unknown and variable.

**Figure 8:**
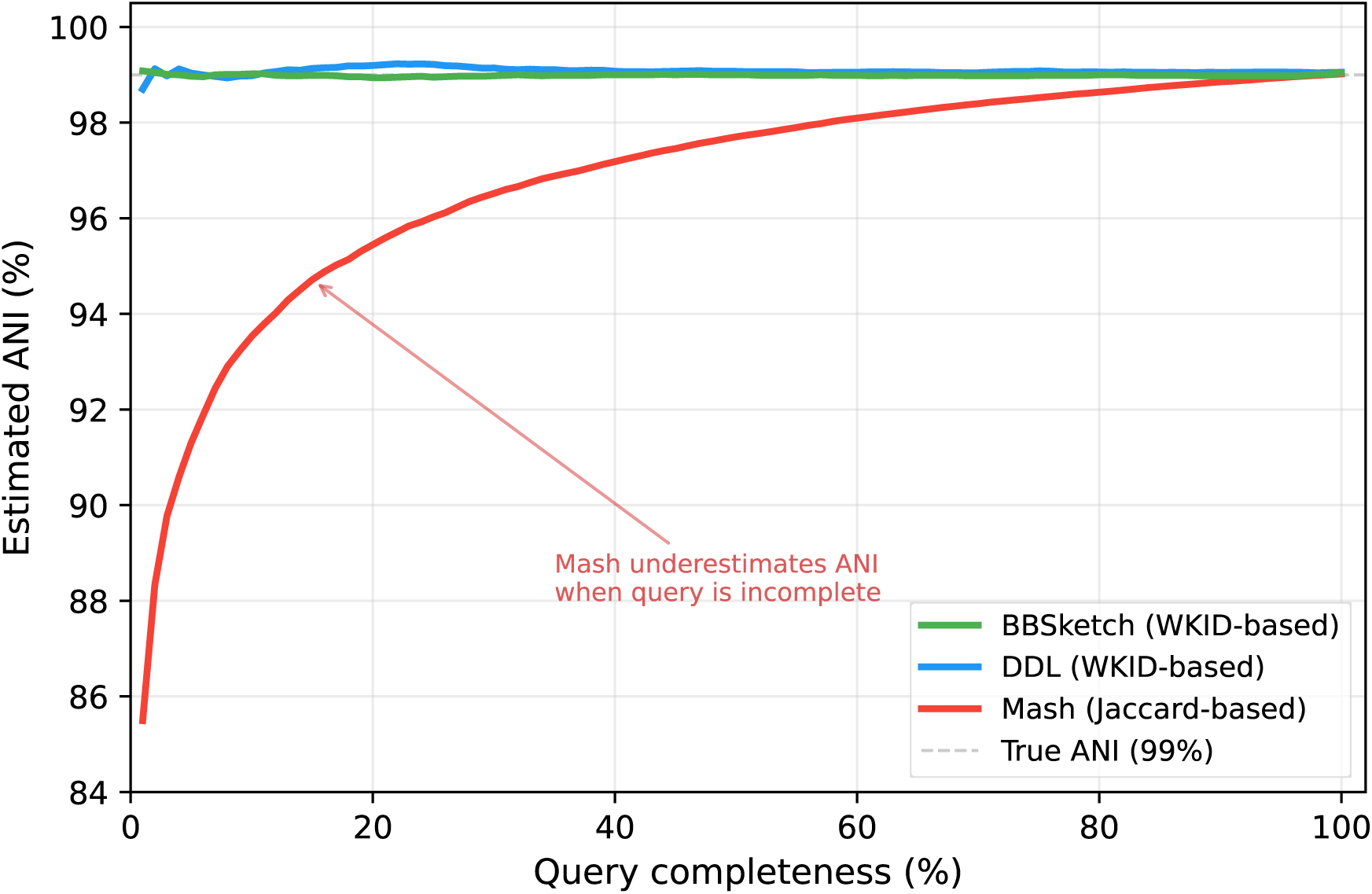
ANI estimation accuracy vs. query genome completeness. *E. coli* K-12 mutant (99% true ANI) truncated to 1–100% of original length and compared against the full reference. DDL and BBSketch (WKID-based) produce stable estimates regardless of query size. Mash (Jaccard-based) underestimates ANI severely when the query is smaller than the reference. B=2,048, k=31.

### 11.3 Size Scaling

The scaling test measures false similarity between a 5M-element reference sketch and query sketches of varying size (2K–64M elements) from independent random streams, averaged over 50 trials per size point.

DDL produces **near-zero false-matching registers** on unrelated random input across the full size range (average 0.5 at 4M elements — consistent with the expected collision rate of ∼0.37 per pair). SetSketch is similar, averaging 0.6 false matches at 4M. LL6 produces hundreds of false matches at every tested size above 4K, peaking at 346 false matches at 4M elements (Figure 9). At 4M elements (average bacterial genome), LL6’s false matches translate to a spurious WKID of 0.318 — equivalent to an ANI estimate of 96.4%, indistinguishable from closely related bacteria. DDL and SetSketch both report WKID ≈ 0 across the full range.

**Figure 9:**
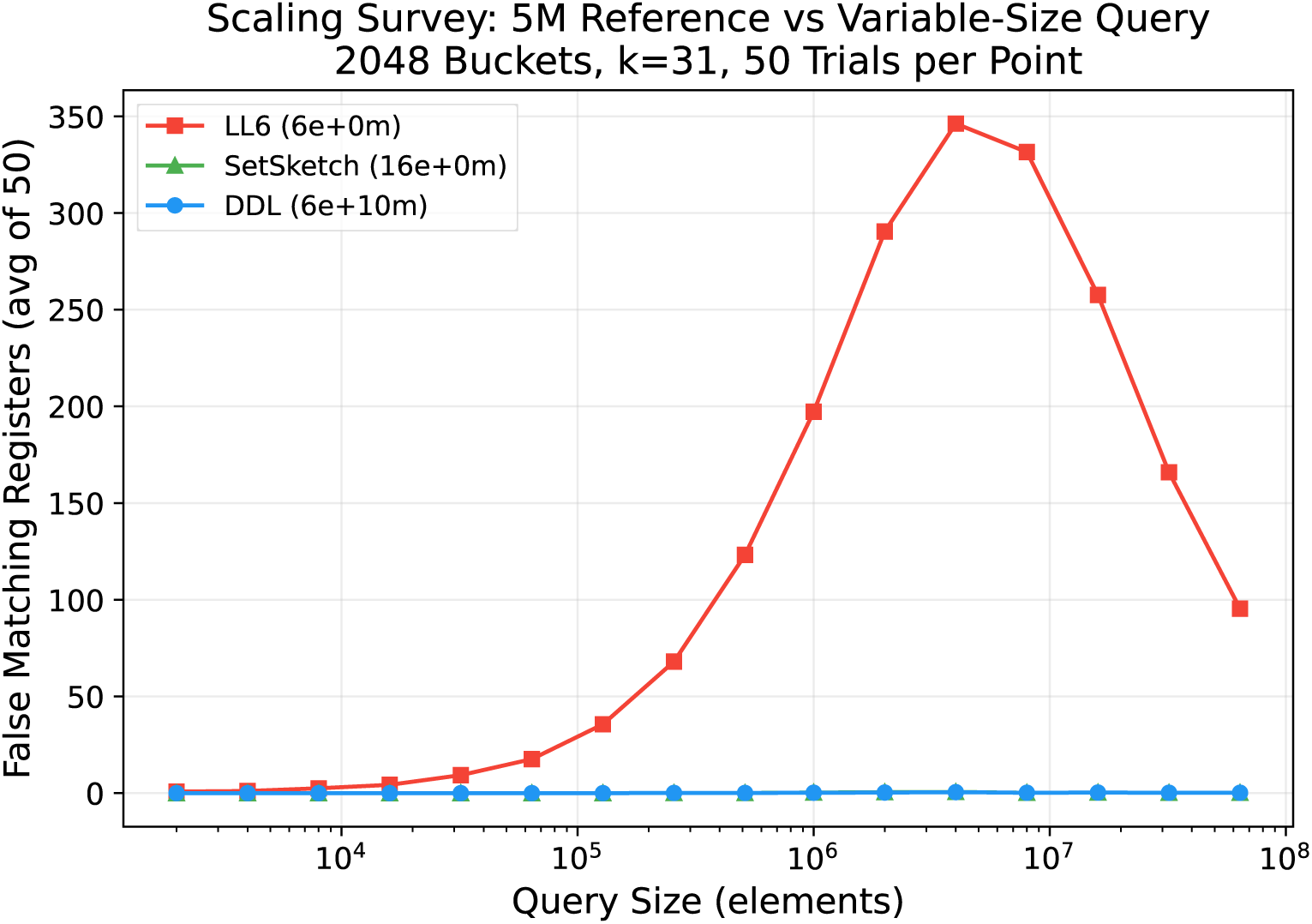
Scaling survey: false matching registers on unrelated random input. 5M-element reference vs variable-size query, 2,048 buckets, 50 trials per point. Elements are random 64-bit hashes; no k-mers or genomes are involved. LL6 peaks at ∼350 false matches near equal size; DDL and SetSketch remain near zero.

### 11.4 Construction Speed

DDL and LL6 have **similar construction speed**; SetSketch is **3.5× slower** due to its per-element logarithm (Figure 10):

**Figure 10:**
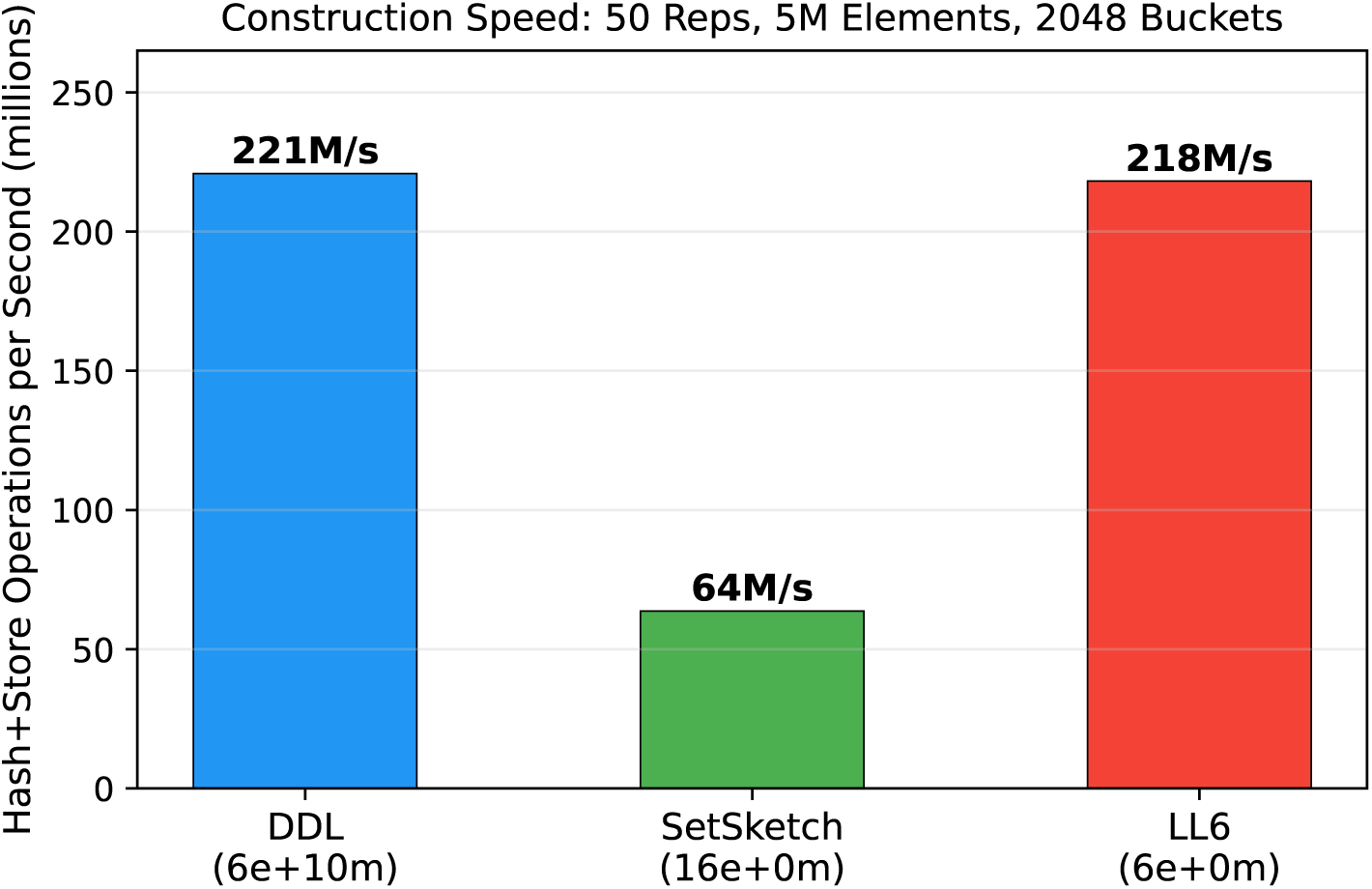
Construction speed comparison. DDL and LL6 are similar — the mantissa adds no measurable overhead. SetSketch is 3.5× slower due to per-element logarithm computation. Single-threaded, 50 reps, 5M elements, 2,048 buckets, fixed-frequency CPU.

The 1.3% difference between DDL and LL6 is within measurement noise. DDL’s mantissa computation (a bit shift and mask, both single-cycle operations) adds negligible overhead to the hash-and-store pipeline, which is dominated by hash computation and memory access. DDL’s early exit mask (Section 3.2) rejects the vast majority of elements before any register access, and the mantissa is computed only for the small fraction that pass this filter. LL6 lacks this optimization, yet construction throughput is identical because both are dominated by hash computation. All speed measurements are single-threaded with a single concurrent estimator on a single core with fixed clock frequency (Section 10.9).

SetSketch’s per-element `Math.log()` call is substantially more expensive than DDL’s bit-shift encoding, though like DDL, it could probably be improved by an early exit. However, the finer logarithm would make dynamically updating the early exit bound more complex and likely expensive.

### 11.5 SIMD Acceleration

DDL’s 16-bit registers pack 16 keys into a 256-bit vector, enabling SIMD-accelerated comparison that 64-bit MinHash keys, using branchy sorted-list merges, cannot exploit. Table 11 quantifies the effect by running DDLCompare with SIMD enabled (default) and disabled (`simd=f`), at 32 threads.

Sketch construction benefits from SIMD only at high thread counts (1.02× at t=1 vs. 2.5× at t=32), because the single-threaded path is dominated by hash computation (which scales well with threads but is not accelerated) rather than parsing (which scales poorly with threads but is SIMD-accelerated). Comparison benefits at all thread counts (2.9× at t=1, 2.5× at t=32), reflecting the direct acceleration of the per-register comparison loop. Preconverting the register arrays from Java’s unsigned char type to signed short arrays allows direct vector loading without per-element type conversion, yielding an additional ∼50% compare speedup (measured at t=1); this workaround would be unnecessary in languages with native uint16 vector support, and is not used in the release for simplicity.

### 11.6 Cardinality Estimation Accuracy

Figure 11 compares cardinality estimation accuracy for DDL and LL6 across the full cardinality range (1 to 16.8M), using 524,288 independent estimators per configuration. Four estimation curves are plotted:

1. **DDL 2048 Hybrid (4 KB)** — DDL’s production estimator (hybridDDL), which blends Linear Counting at low cardinality with mantissa-aware MeanM at high cardinality.
2. **LL6 2048 Hybrid (2 KB)** — LL6’s production estimator (hybridDLL), using the same LC-to-Mean blend but without mantissa information.
3. **LL6 2048 HLL (2 KB)** — raw HyperLogLog estimator at 2,048 buckets, showing the characteristic LC-to-HLL transition spike.
4. **LL6 4096 HLL (4 KB)** — raw HyperLogLog at 4,096 buckets (same memory as DDL), with lower steady-state error but the same transition spike.

**Figure 11:**
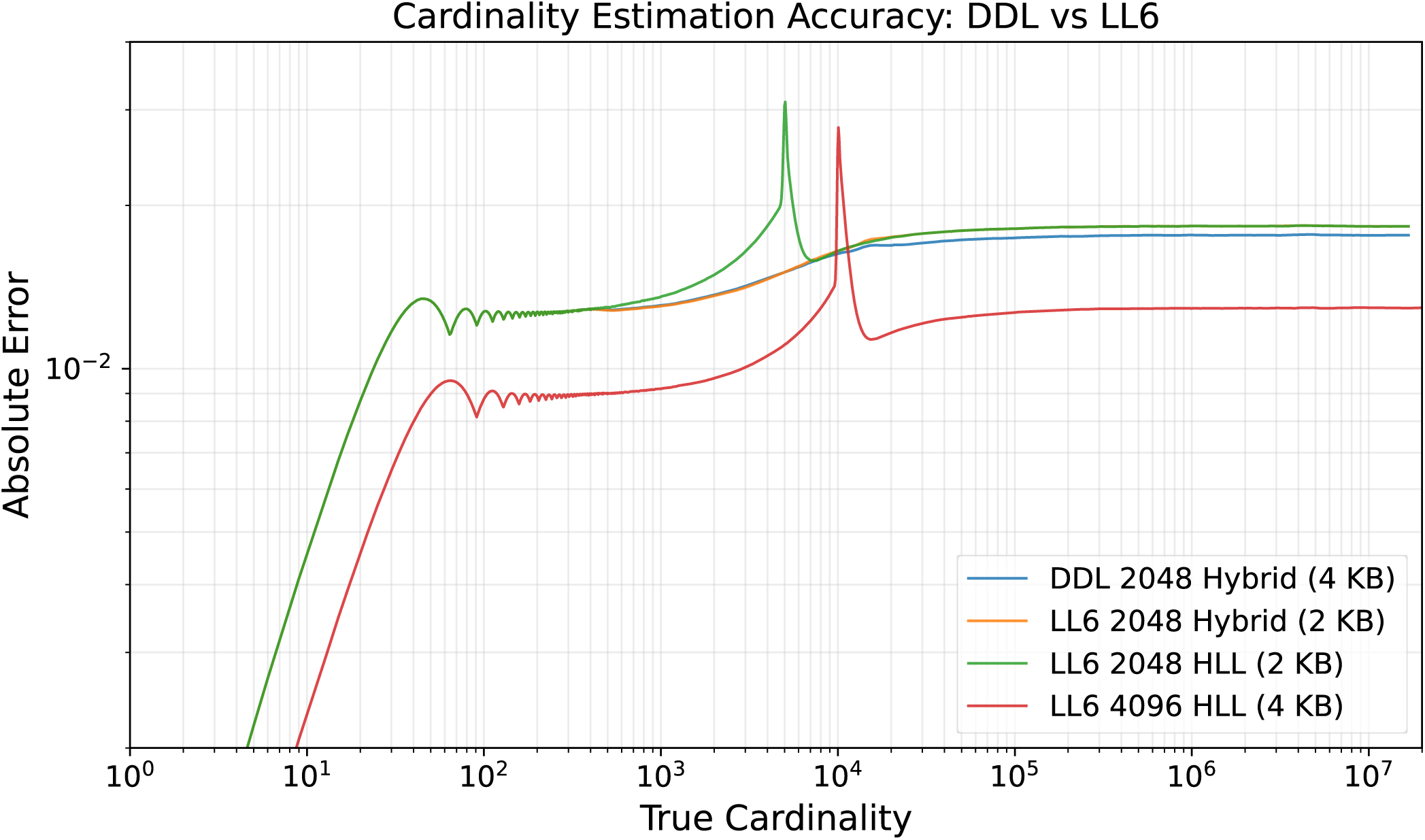
Cardinality estimation accuracy. Log-log plot of absolute error vs. true cardinality. DDL 2048 Hybrid achieves approximately 3.8% relatively lower error than LL6 2048 Hybrid using the mantissa for sub-NLZ accuracy; LL6 4096 HLL achieves lower steady-state error at equal memory but retains the LC-to-HLL transition spike. 524,288 independent estimators per configuration, correction factors enabled.

At equal bucket count (2,048), DDL’s Hybrid is marginally more accurate than LL6’s Hybrid (1.76% vs. 1.83% width-weighted error) — the mantissa provides a small cardinality improvement through MeanM’s finer-grained hash magnitude recovery. Both Hybrid curves eliminate the low-cardinality transition error spike visible in standard HLL.

At equal memory (4 KB), LL6 with 4,096 buckets achieves lower steady-state error than DDL with 2,048 buckets (1.29% vs. 1.76%) — doubling the number of independent observations outweighs the mantissa’s per-register precision. However, despite doubling the bucket count, it still displays much higher peak error.

DDL’s cardinality accuracy is thus similar to LL6 — the mantissa bits neither help nor hurt significantly for cardinality estimation. DDL’s value proposition is not improved cardinality but the additional capabilities (comparison, frequency, index) obtained at no accuracy cost.

### 11.7 Database Query Performance

The three tools were benchmarked using their respective RefSeq reference databases as described in Section 10.6. Sketch construction speed was measured on 14.7 million reads (2,623 Mbp) to ensure timing was dominated by hashing rather than JVM startup.

#### Sketch construction

Sketch speed was measured on 14.7 million reads (2,623 Mbp). DDL constructs a query sketch at 126 Mbp/s (single-threaded) — 6.3× faster than CompareSketch (20 Mbp/s) and 3.4× faster than Mash (37 Mbp/s). DDL’s speed advantage comes from the early exit mask (Section 3.2), which rejects most k-mers before any memory access. CompareSketch’s lower throughput is partially explained by dual k-mer processing (k=32 and k=24 simultaneously). At 32 threads, DDL achieves 1,076 Mbp/s (8.5× speedup), CompareSketch achieves 460 Mbp/s (23× speedup), and Mash does not parallelize sketch construction (37 Mbp/s at both t=1 and p=32). DDL at 32 threads is 29× faster than Mash and 2.3× faster than CompareSketch.

#### Reference loading

DDL loads 200,687 sketches (411M 16-bit keys) via DDLCompare’s multithreaded loader (DDLLoaderMT). Loading throughput scales well with threads (Tables 13–14). Mash’s lower loading time reflects its memory-mapped binary format.

**Table 4:** Average number of k-mers shared between random pairs, grouped by the taxonomic level of the pair’s lowest common ancestor. RefSeq genome database, 500,000 random pairs, k=31, B=2,048, exponent=5. Noise = weighted average at class+ distance. *Unmerged*: 200,687 sketches, with organelles (mitochondria, plastids) and plasmids kept as separate records. *Merged*: 159,567 sketches, with organelles and plasmids combined into their host genome sketches by TaxID.

| Database | Genus | Family | Order | Class | Phylum | Kingdom | Noise |
| --- | --- | --- | --- | --- | --- | --- | --- |
| Unmerged, no blacklist | 162.73 | 74.68 | 19.68 | 32.12 | 3.54 | 1.28 | 11.51 |
| Unmerged, blacklisted (23,020) | 160.37 | 53.25 | 6.17 | 2.33 | 0.23 | 0.16 | 0.86 |
| Merged, no blacklist | 182.65 | 35.13 | 11.30 | 15.49 | 4.11 | 1.60 | 5.98 |
| Merged, blacklisted (23,020) | 181.39 | 26.35 | 2.74 | 1.22 | 0.25 | 0.17 | 0.47 |

**Table 5:** Average shared keys and noise floor for the k=25, B=2,048, exponent=5 RefSeq database, 2,000,000 random pairs. Noise = average shared keys at class+ distance.

| Database | Species | Genus | Family | Order | Class | Phylum | Kingdom | Noise |
| --- | --- | --- | --- | --- | --- | --- | --- | --- |
| Unmerged, no blacklist | 1251.56 | 182.89 | 86.17 | 24.21 | 42.49 | 5.04 | 1.71 | 2.61 |
| Unmerged, blacklisted<br>(1,651) | 1251.42 | 181.86 | 78.79 | 17.28 | 20.89 | 1.18 | 0.32 | 1.21 |
| Merged, blacklisted (1,651) | 1352.60 | 198.62 | 43.13 | 9.55 | 9.88 | 1.35 | 0.36 | 0.62 |

**Table 6:**
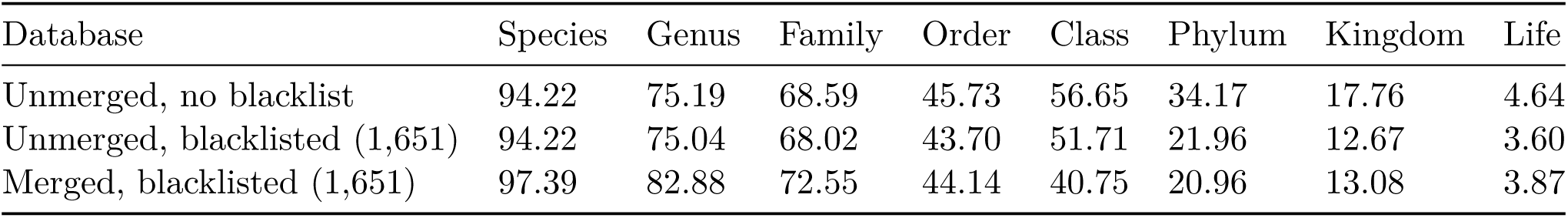
Average estimated ANI (WKID^1^*^/k^*, percent) for the same configurations. Species- and genus-level ANI is essentially unchanged by the blacklist, confirming signal preservation.

**Table 7:**
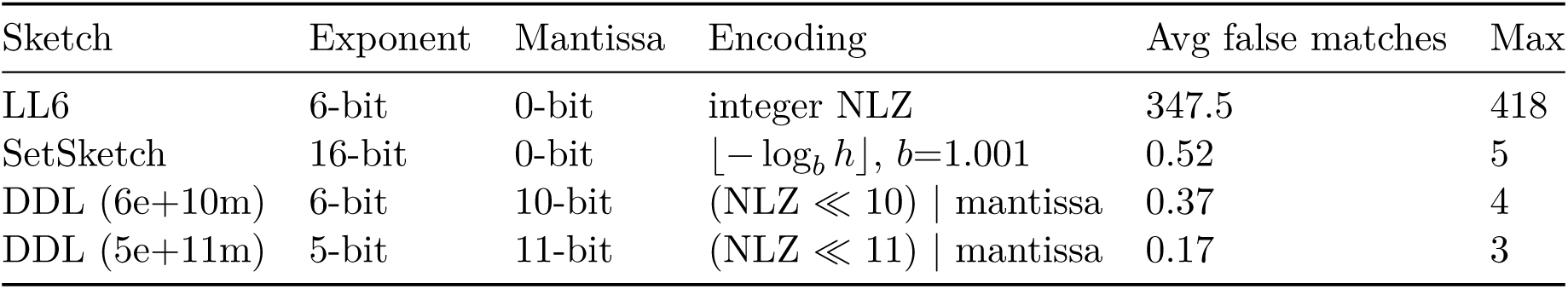
Collision rate on unrelated random input. 100 sketches, 4,950 pairs, 4M elements each (Xoshiro256 PRNG), 2,048 buckets, all 16-bit registers.

**Table 8:** ANI estimation accuracy averaged over 50 independent trials per level. 1 MB random genome, mutate.sh uniform SNPs, quantumaligner.sh measured identity, 2,048 buckets, k=25, exponent=5. *σ* = standard deviation across trials.

| Design ANI | Aligned ANI | DDL ANI | LL6 ANI | Aligned $\sigma$ | DDL $\sigma$ | DDL matches |
| --- | --- | --- | --- | --- | --- | --- |
| 99.9% | 99.90% | 99.91% | 99.94% | 0.003% | 0.012% | 1,951 |
| 99.5% | 99.50% | 99.52% | 99.66% | 0.007% | 0.026% | 1,618 |
| 99.0% | 98.99% | 99.02% | 99.33% | 0.009% | 0.041% | 1,300 |
| 98.0% | 98.00% | 98.04% | 98.70% | 0.012% | 0.070% | 884 |
| 97.0% | 97.00% | 97.06% | 98.15% | 0.018% | 0.091% | 627 |
| 95.0% | 95.01% | 95.03% | 97.19% | 0.022% | 0.173% | 328 |
| 90.0% | 90.05% | 90.08% | 95.84% | 0.031% | 0.422% | 77.1 |
| 87.0% | 87.11% | 87.02% | 95.48% | 0.030% | 0.573% | 32.0 |
| 85.0% | 85.18% | 84.80% | 95.38% | 0.034% | 0.777% | 16.8 |
| 80.0% | 80.46% | 78.34% | 95.28% | 0.036% | 11.329% | 4.2 |
| 79.0% | 79.53% | 77.68% | 95.25% | 0.037% | 11.249% | 3.4 |
| 77.0% | 77.71% | 66.97% | 95.24% | 0.037% | 27.071% | 1.9 |

**Table 9:** Construction speed. Single-threaded, 50 reps, 5M random hash elements, 2,048 buckets, fixed-frequency CPU.

| Sketch | Adds/sec | 50 reps, 5M elements |
| --- | --- | --- |
| DDL (6e+10m) | 220,862,440 | 1.13s total |
| SetSketch (16e+0m) | 63,703,737 | 3.92s total |
| LL6 (6e+0m) | 218,106,755 | 1.15s total |

**Table 10:** SIMD effectiveness at 32 threads (wall time). Sketch speed reflects SIMD-accelerated fastq parsing; compare speed reflects SIMD-accelerated 16-way register comparison via `jdk.incubator.vector`.

| Phase | SIMD on | SIMD off | Speedup |
| --- | --- | --- | --- |
| Sketch query (4,832 Mbp) | 4.1 s | 10.4 s | 2.5× |
| Brute force compare (1,000 × 200,687) | 6.7 s | 16.9 s | 2.5× |

**Table 11:**
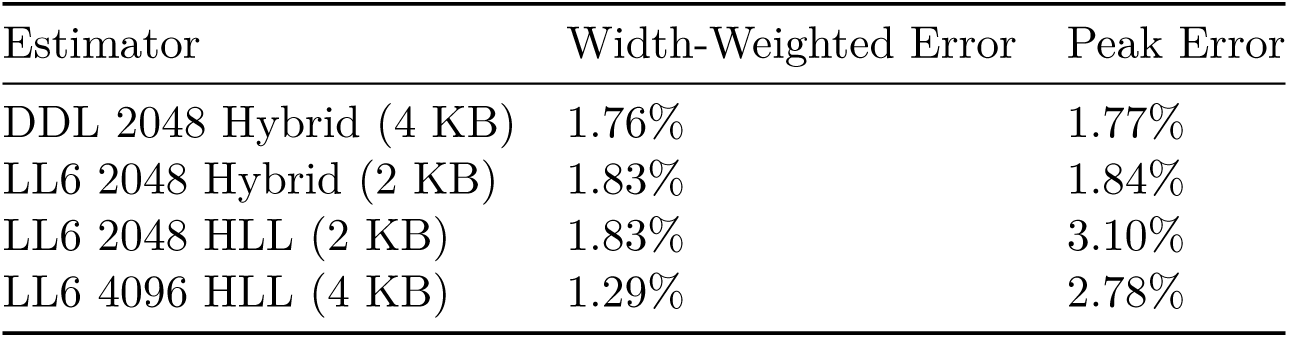
Cardinality estimation accuracy. 524,288 estimators, correction factors enabled, cardinality 1 to *B* × 8,192.

**Table 12:** Single-threaded performance (t/p = 1). Reference databases: DDL 200,687 sketches (411M 16-bit keys); CompareSketch 85,155 sketches (1.21B 64-bit keys); Mash 91,282 sketches (91.3M 64-bit keys). MKeys/s denominators use total reference keys. Queries/s = complete database searches per second, amortized over 1,000–10,000 queries. Index efficiency = 1 - (indexed comparisons / brute-force comparisons). ^†^Mash uses memory-mapped binary I/O; value reflects file size per key, not Java heap.

| Phase | DDLCompare | CompareSketch | Mash |
| --- | --- | --- | --- |
| Sketch query (Mbp/s) | 126 | 20 | 37 |
| Ref load (MKeys/s) | 19.5 | 16 | 43 |
| Brute force (MKeys/s) | 2,260 | 120 | 72 |
| Brute force (queries/s) | 5.5 | 0.21 | 0.78 |
| Index build (MKeys/s) | 5.4 | 3.1 | — |
| Index query (queries/s) | 260 | 2.1 | — |
| Index efficiency | 96.3% | — | — |
| Heap, no index (B/key) | 2.4 | 11.6 | 7.9<sup>†</sup> |
| Heap, indexed (B/key) | 9.7 | 25.6 | — |

**Table 13:**
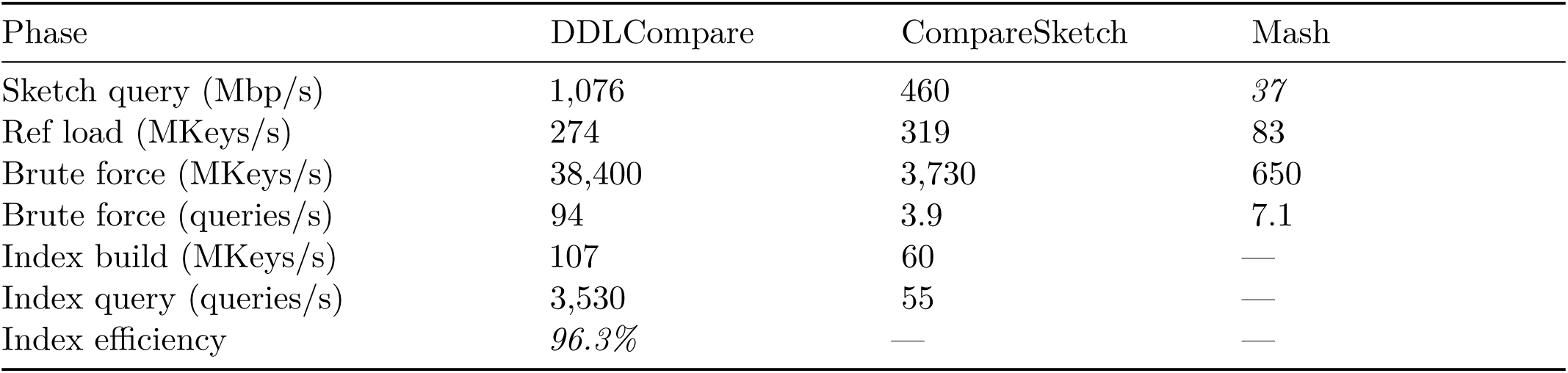
32-threaded performance (t/p = 32). *Italic* values do not parallelize and are unchanged from single-threaded. DDL ref loading parallelizes via QuickClade; Mash sketch construction does not parallelize. Indexed queries used a batch of 10,000 queries; brute-force queries/s used 1,000 (t=1) or 10,000 (t=32). ^†^Mash uses memory-mapped binary I/O.

| Phase | DDLCompare | CompareSketch | Mash |
| --- | --- | --- | --- |
| Sketch query (Mbp/s) | 1,076 | 460 | *37* |
| Ref load (MKeys/s) | 274 | 319 | 83 |
| Brute force (MKeys/s) | 38,400 | 3,730 | 650 |
| Brute force (queries/s) | 94 | 3.9 | 7.1 |
| Index build (MKeys/s) | 107 | 60 | — |
| Index query (queries/s) | 3,530 | 55 | — |
| Index efficiency | *96.3%* | — | — |

**Table 14:** All-to-all comparison at 128 threads (64 cores, single-socket AMD EPYC 9555P). Each tool compares its native database against itself; queries/s = sketches ÷ wall-clock seconds; MKeys/s = total reference keys × queries/s (brute force only; indexed rows omitted because the index avoids most key comparisons). DDL: 200,687 sketches (411M 16-bit keys); CompareSketch: 85,155 sketches (1.21B 64-bit keys); Mash: 91,282 sketches (91.3M 64-bit keys).

| Tool | Mode | Sketches | Time (s) | Queries/s | MKeys/s |
| --- | --- | --- | --- | --- | --- |
| DDL | indexed | 200,687 | 30.4 | 6,601 | — |
| DDL | brute force | 200,687 | 363.0 | 553 | 227,300 |
| CompareSketch | indexed | 85,155 | 650.8 | 131 | — |
| CompareSketch | brute force | 85,155 | 5,638.8 | 15.1 | 18,300 |
| Mash | brute force | 91,282 | 5,186.7 | 17.6 | 1,610 |

#### Brute force comparison

DDL compares 2,260 MKeys/s single-threaded and 38,400 MKeys/s at 32 threads (17× scaling), via SIMD-accelerated branchless comparison of 16-bit registers (16 pairs per AVX-256 instruction). CompareSketch achieves 120 MKeys/s single-threaded and 3,730 MKeys/s at 32 threads. Mash achieves 72 MKeys/s single-threaded and 650 MKeys/s at 32 threads via sorted-merge of 64-bit hash lists. In terms of complete database queries, DDL performs 5.5 brute force queries/s at t=1 and 94 queries/s at t=32 against 200,687 references, compared to 0.21 and 3.9 queries/s for CompareSketch (85,155 references) and 0.78 and 7.1 queries/s for Mash (91,282 references).

#### Inverted index

DDL’s inverted index identifies candidate references sharing ≥ 5 matching registers (one below the reliability floor of 6, to avoid missing borderline true positives from small genome fragments), then performs full pairwise comparison on those candidates to compute WKID and ANI. The index eliminates 96.3% of comparisons (index efficiency), reducing 200,687 brute-force comparisons per query to ∼7,400 candidate comparisons. With this filtering, DDL completes 260 indexed queries per second single-threaded and 3,530 queries/s at 32 threads against 200,687 references (amortized over 10,000 queries) — 3.8 ms and 0.28 ms per query, respectively. The index builds in 76 seconds single-threaded or 3.8 seconds at 32 threads, with 36.4M populated cells out of 134.2M total (27% populated, 73% null). CompareSketch’s KmerTableSet index completes 2.1 queries/s single-threaded or 55 queries/s at 32 threads (amortized over 100–1,000 queries); CompareSketch’s filtered comparisons (∼1,900 per query) parallelize effectively. Mash does not provide an inverted index.

#### All-to-all comparison

To measure end-to-end performance on a realistic workload, each tool compared its full RefSeq database against itself at 128 threads (64 cores). DDL’s indexed search of 200,687 sketches completes in 30 seconds — over 375× faster per query than Mash’s brute-force search of 91,282 sketches, and over 50× faster than CompareSketch’s index, while searching a 2.2× larger database. In raw key comparison throughput, DDL brute force achieves 227,300 MKeys/s — 12× faster than CompareSketch (18,300 MKeys/s) and 141× faster than Mash (1,610 MKeys/s).

#### Memory

DDL requires 2.4 bytes per key without indexing (1 GB for 411M keys) and 9.7 bytes per key with index (4 GB). CompareSketch requires 11.6 bytes per key without indexing (14 GB for 1.21B keys) and 25.6 bytes per key with index (31 GB). DDL loads 2.4× more sketches in 14× less memory without index, or 7.8× less memory with index. DDL’s inverted index (a sparse 2D array bounded by the 65,536-value register space) is far more compact than CompareSketch’s KmerTableSet (508 million 64-bit hashcodes). Mash uses memory-mapped binary I/O at 7.9 bytes per key (720 MB file for 91.3M keys), which is not directly comparable to Java heap measurements.

### 11.8 Completeness Estimation

DDL’s completeness metric (Section 5.3) estimates the fraction of a reference genome represented in a query. To validate accuracy, the *E. coli* K-12 genome (4.64 Mbp) was truncated to 5–100% of its length (in 5% increments) and compared against the full reference at four ANI levels: 100% (identical sequence), 99% (1% substitution via `mutate.sh`), 95% (5% substitution), and 90% (10% substitution).

Figure 12 shows that DDL completeness tracks actual completeness closely at all four ANI levels. At 100% ANI, the estimate falls on the diagonal — a 50% truncation yields a completeness estimate of 0.494. At 99% ANI, the estimate remains accurate across the full range (maximum deviation <6%). At 95% ANI, the estimate shows more noise at low completeness but remains well-calibrated overall. Even at 90% ANI (k-mer survival probability 0.90^25^ ≈ 7.2%, approximately 147 expected matching registers at full completeness), the estimate tracks the diagonal without systematic bias — a significant advantage of k=25 over longer k-mer lengths, where the sparser signal causes systematic overshoot.

**Figure 12:**
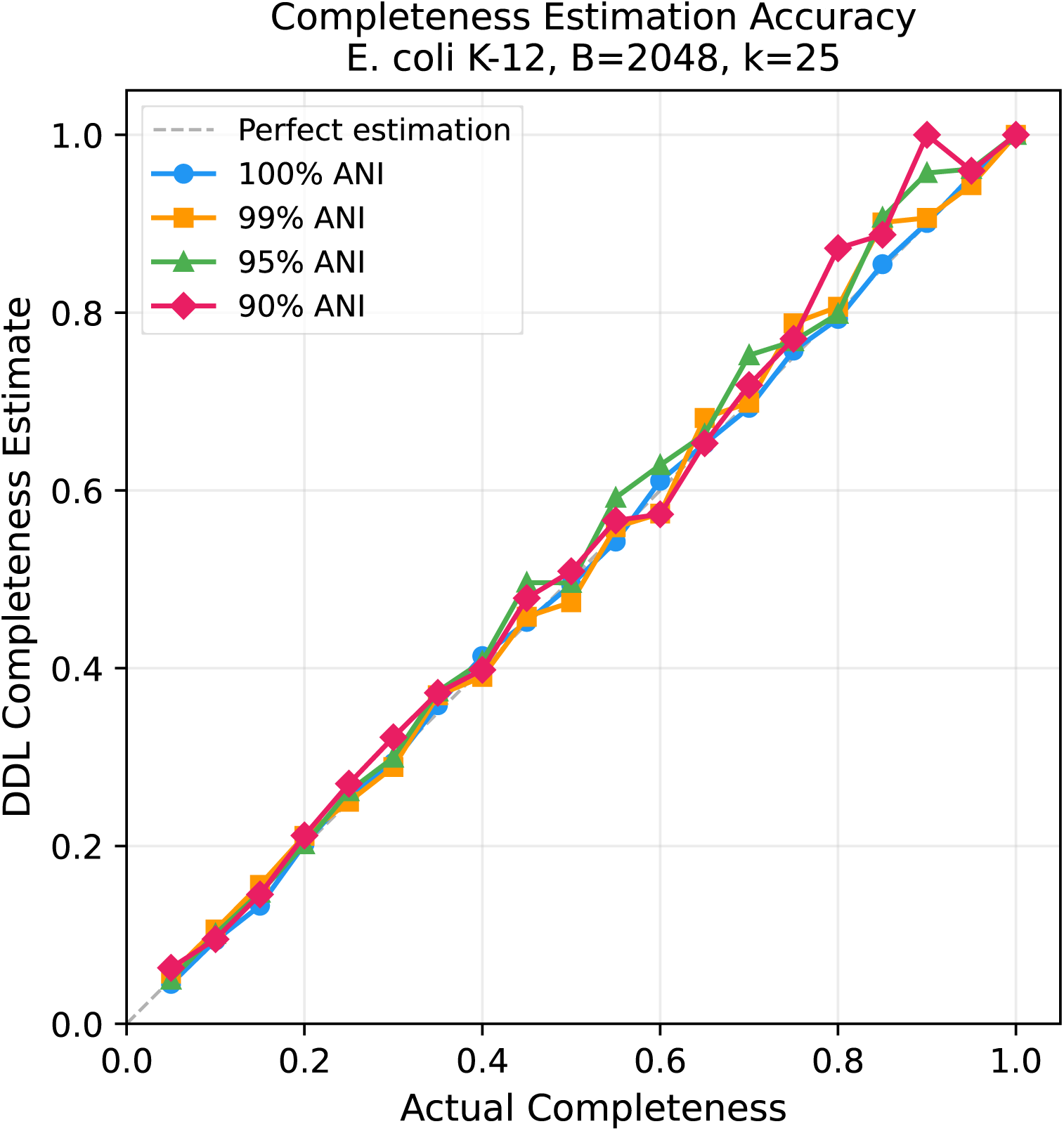
Completeness estimation accuracy. DDL completeness estimate vs. actual genome fraction at 100%, 99%, 95%, and 90% ANI. *E. coli* K-12 truncated to 5-100% of its length, compared against the full reference. All four ANI levels track the diagonal closely. Dashed line = perfect estimation. B=2,048, k=25.

### 11.9 Genome Size Estimation

DDL’s per-register frequency counter (Section 7) enables genome size estimation from raw sequencing reads — a task normally requiring exact k-mer counting with memory proportional to the number of distinct k-mers. The k-mer frequency histogram produced by DDL’s `loglog.sh` tool can be analyzed by a peak-finding algorithm that separates error k-mers (low depth, appearing once or twice due to sequencing errors) from genomic k-mers (clustered around the true coverage depth), estimating genome size from the area under the genomic peak.

Figure 13 demonstrates this on a real sequencing library of *Micromonospora rosaria* (assembled genome size 7.28 Mbp, 17× coverage). DDL with 262,144 buckets (512 KB sketch) reports a cardinality of 21.6 million distinct k-mers — 3× the actual genome size, inflated by 14.3 million error k-mers (66% of total). Despite this, the peak-finding algorithm correctly identifies the main coverage peak at depth 16 and estimates the genome size at 7,449,225 — within 2.3% of the reference value.

**Figure 13:**
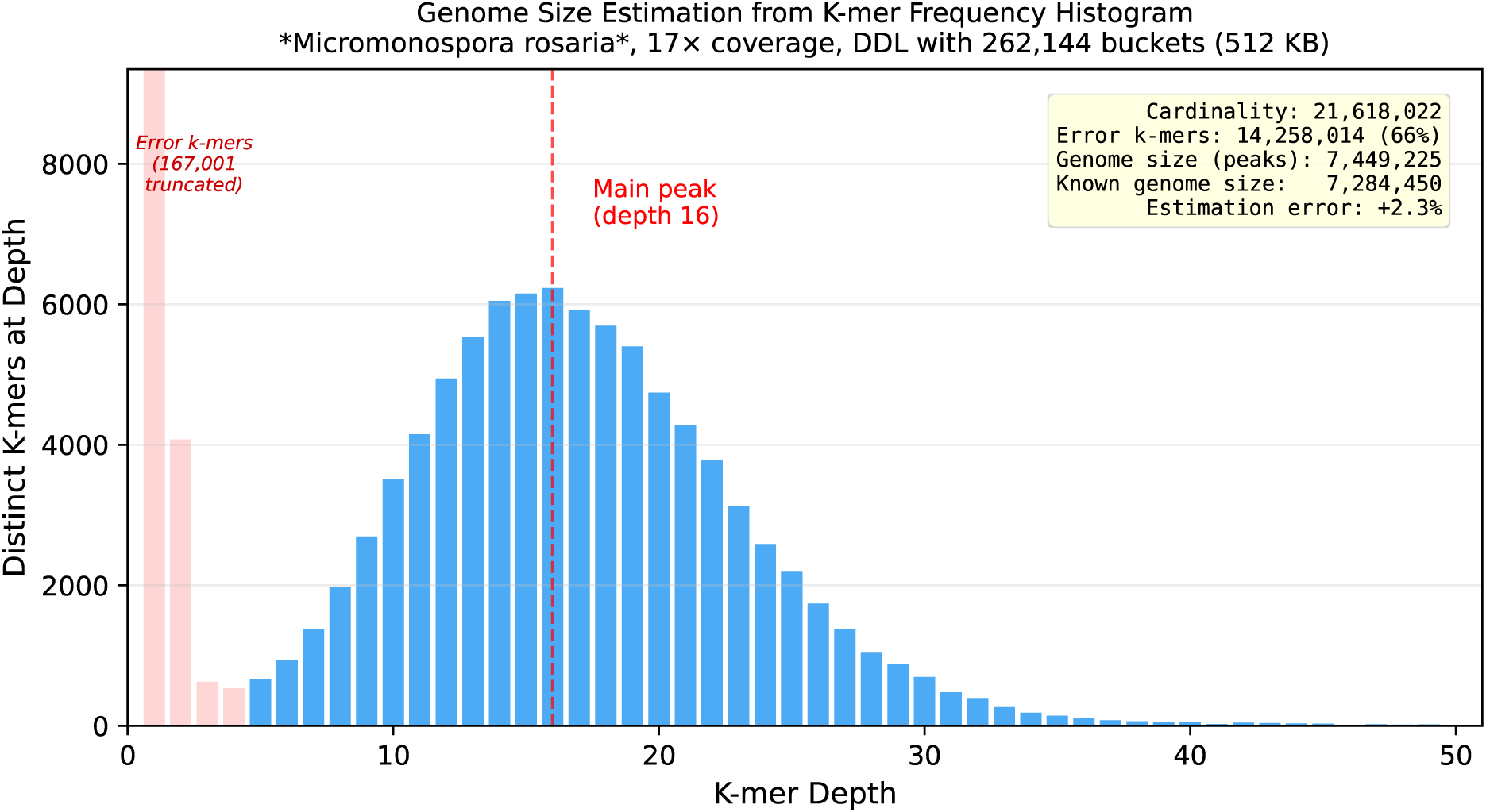
Genome size estimation from a real sequencing library. *Micromonospora rosaria*, 17× coverage, k=31, DDL with 262,144 buckets (512 KB). The depth-1 bar (167,001 buckets) is truncated for readability. All bar heights represent bucket counts; multiply by cardinality/buckets (≈82.5×) for estimated true k-mer counts. Despite cardinality being 3× the genome size due to 14.3 million error k-mers (66% of total), the peak-finding algorithm estimates genome size within 2.3% of the known 7.28 Mbp.

### Comparison with exact and approximate counting

Three tools were timed on the same *M. rosaria* library (178 MB compressed FASTQ, Dori cluster):

**Table 15:**
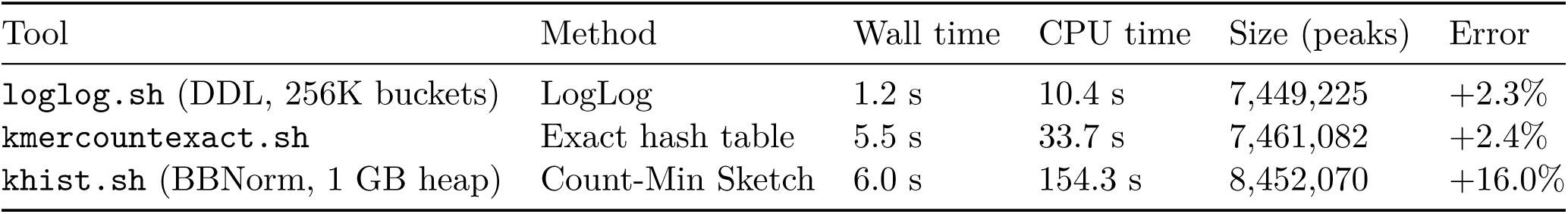
Genome size estimation speed and accuracy. Known genome size: 7,284,450 bp.

DDL matches exact counting’s accuracy (+2.3% vs +2.4%) while running 4.5× faster in wall time and 3.2× faster in CPU time. The Count-Min Sketch approach (`khist.sh`) uses comparable wall time to exact counting but uses 4.6× as much CPU time (multithreaded), and overestimates genome size by 16% because hash collisions in the CMS inflate k-mer counts and blur the coverage peak. KmerCountExact identifies the main peak at depth 18 (the gold standard); DDL identifies it at depth 16 (close, but slightly shifted due to histogram resolution); CMS identifies it at depth 14 (substantially shifted by collision-inflated counts). The genome size estimate depends on the area under the peak rather than peak position alone, which is why DDL’s +2.3% accuracy closely matches KmerCountExact’s +2.4% despite the slight peak shift.

The accuracy of genome size estimation also depends on histogram resolution, which is determined by bucket count. Figure 14 shows this scaling: at 2,048 buckets (4 KB), the estimate is 11% high; at 32,768 buckets (64 KB), it converges to within 7%; and at 262,144 buckets (512 KB), it is within 2.3%. KmerCountExact (exact counting, unbounded memory) achieves 2.4% on the same data — DDL matches this accuracy from a 512 KB sketch.

**Figure 14:**
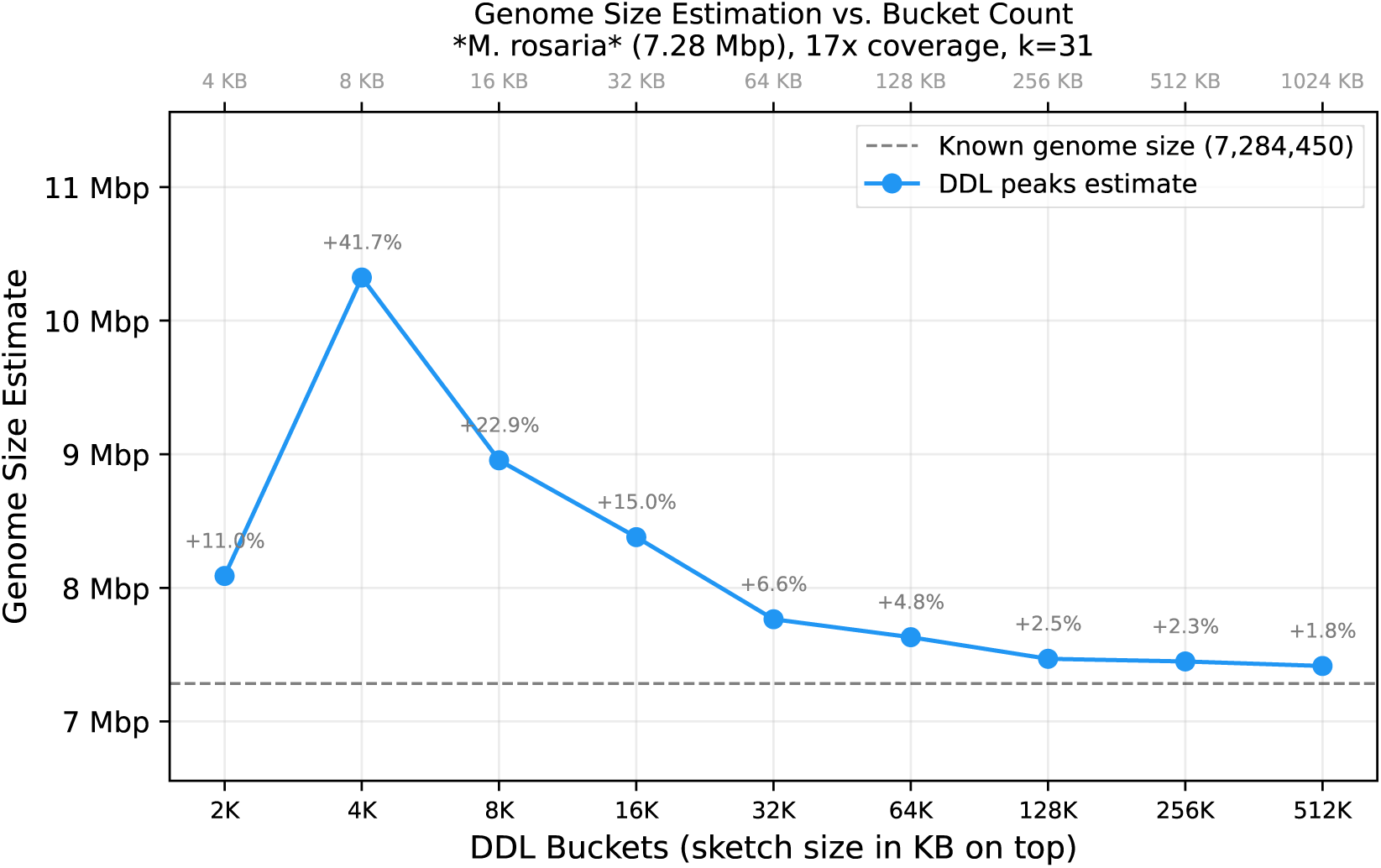
Genome size estimation accuracy vs. DDL bucket count. *M. rosaria* (known genome 7.28 Mbp), 17× coverage, k=31. Sketch size (top axis) ranges from 4 KB to 1 MB. The estimate converges to within 2.3% of the known genome size at 262K buckets, matching KmerCountExact’s 2.4% accuracy.

## 12. Discussion

### 12.1 When to Use DDL vs. Alternatives

#### DDL vs. LogLog

Use DDL when you need *any* comparison capability in addition to cardinality. Use pure cardinality-optimized LogLog variants such as AVLL (Bushnell 2026a) when memory is the binding constraint and comparison is not needed (e.g., embedded systems, per-flow counters in network hardware).

#### DDL vs. MinHash

BBSketch (BBTools’ MinHash implementation) and its server-based companion SendSketch provide a mature ecosystem for taxonomic identification via MinHash sketches. SendSketch sends a MinHash sketch to a remote server hosting a reference database, while CompareSketch performs local comparison. DDL does not replace this ecosystem, but complements it: QuickClade (BBTools) uses DDL sketches internally to complement its short-kmer-frequency Clade index, allowing dual lookup from orthogonal metrics to calculate a top hit LCA and thus accurately estimate taxonomic level assignment confidence. Use MinHash/SendSketch when you need the full taxonomic ecosystem (e.g., remote servers, community databases); use DDL when you need faster local comparison, cardinality estimation, or memory-constrained deployment (DDL uses 4× less memory than MinHash at equal sketch size).

#### DDL vs. Bloom Filter/Exact Counts

DDL dramatically outperforms these data structures in both time and memory, when the desired output is an element frequency histogram or composition histogram. It does not support set inclusion or element count queries, so explicit hashtables or Bloom Filters / Count-Min Sketches are still necessary for those operations.

### 12.2 The Inverted Index as Killer Application

For database search at scale, the inverted index (Section 6) may be DDL’s most significant contribution. A database of *N* sketches can be searched in O(*B* + *M*) time per query, where *M* is the total number of matching index entries. When most database sketches are unrelated to the query, *M* is small and query time is dominated by the O(*B*) scan, which grows with sketch size rather than database size — ∼40× faster than exhaustive pairwise comparison at *N* = 200,687 (Section 11.7), with the gap widening as *N* increases.

The index is possible because DDL’s 65,536 possible register values hit the practical sweet spot for indexing — too few values (LL6) fails due to high collision rates and too many (MinHash) requires a keyset, as discussed in Section 6.3.

### 12.3 Limitations

- **Practical limits of discrete sampling.** DDL’s reliability drops at high sequence divergence or extremely low completeness because a shared k-mer must win its respective bucket in *both* sketches to be detected (Section 8.3, Section 11.2). While this detection floor can be extended by adjusting k-mer length or bucket count, DDL is inherently constrained by this discrete-sampling threshold.
- **Hash-bound construction.** DDL’s 16-bit registers allow SIMD-accelerated comparison, but sketch construction is dominated by scalar hash computation and does not currently benefit from SIMD vectorization.
- **Approximate frequency tracking.** The per-register observation counter (Section 7) records only the winning element’s multiplicity. It characterizes the overall frequency distribution of a stream but cannot answer point queries for specific elements.

### 12.4 Future Work

#### Long-read assembly overlap detection

DDL’s inverted index could extend to the overlap detection phase of overlap-layout-consensus (OLC) assembly, where each read is compared against every other read. Sketching each read into a DDL and using the inverted index for candidate identification would provide O(*B* + *M*) overlap detection per query. Replacing the per-register counter with a *position array* recording k-mer position rather than frequency could enable overlap geometry estimation directly from the sketch comparison.

#### Counter arrays as stream signatures

DDL’s per-register frequency counters (Section 7.1) could serve as fixed-size signatures for distributional similarity between streams via cosine similarity, measuring whether two streams have similar frequency distributions regardless of shared elements. Combined with per-register GC content (Section 7.4), this provides a two-dimensional stream fingerprint. Validation in domains such as network traffic classification or content fingerprinting is left for future work.

## 13. Conclusion

DynamicDemiLog extends the LogLog register with a floating-point encoding — an integer NLZ exponent plus a fractional inverted mantissa — that preserves all existing cardinality estimation machinery while enabling meaningful per-register comparison, frequency tracking, and inverted index construction.

On synthetic input, DDL achieves a 930× reduction in false register matches compared to standard 6-bit LogLog (Section 11.1). On real nucleotide sequences with controlled mutation, DDL accurately estimates ANI via WKID^1^*^/k^* down to approximately 79% identity (at k=25, 5e+11m), while LL6 reports approximately 95% ANI on unrelated or distantly related inputs. DDL achieves this at similar construction speed and produces near-zero spurious similarity on unrelated inputs.

The mantissa is free: computed from bits already in the hash, requiring only a shift and a mask. Any system using HyperLogLog for cardinality gains similarity, containment, frequency, and index support by switching to DDL with no performance penalty and a ∼2–7× increase in memory per bucket.

DDL is implemented in Java as part of the BBTools suite and is available as open-source software.

## Acknowledgements

Simulations and benchmarks were conducted on the Dori computing cluster at the Joint Genome Institute, Lawrence Berkeley National Laboratory. The work conducted by the U.S. Department of Energy Joint Genome Institute (https://ror.org/04xm1d337), a DOE Office of Science User Facility, is supported by the Office of Science of the U.S. Department of Energy operated under Contract No. DE-AC02-05CH11231.

The author thanks Noire for help writing and generating data for this project.

## Disclosure of AI assistance

This work made extensive use of large language model (LLM) AI tools, including Anthropic Claude and Google Gemini. The initial draft of this manuscript was prepared with AI assistance; Gemini provided detailed editorial feedback and suggestions. All algorithmic design decisions, experimental methodology, interpretation of results, and scientific conclusions are solely those of the author.

